# Whole-genome resequencing data support a single introduction of the invasive white pine sawfly, *Diprion similis*

**DOI:** 10.1101/2022.09.14.507957

**Authors:** Jeremy S. Davis, Sheina Sim, Scott Geib, Brian Sheffler, Catherine R. Linnen

## Abstract

Biological introductions are unintended “natural experiments” that provide unique insights into evolutionary processes. Invasive phytophagous insects are of particular interest to evolutionary biologists studying adaptation, as introductions often require rapid adaptation to novel host plants. However, adaptive potential of invasive populations may be limited by reduced genetic diversity—a problem known as the “genetic paradox of invasions”. One potential solution to this paradox is if there are multiple invasive waves that bolster genetic variation in invasive populations. Evaluating this hypothesis requires characterizing genetic variation and population structure in the introduced range. To this end, we assemble a reference genome and describe patterns of genetic variation in the introduced white pine sawfly, *Diprion similis*. This species was introduced to North America in 1914, where it has undergone a rapid host shift to the thin-needled eastern white pine (*Pinus strobus*), making it an ideal invasion system for studying adaptation to novel environments. To evaluate evidence of multiple introductions, we generated whole-genome resequencing data for 64 *D. similis* females sampled across the North American range. Both model-based and model-free clustering analyses supported a single population for North American *D. similis*. Within this population, we found evidence of isolation-by-distance and a pattern of declining heterozygosity with distance from the hypothesized introduction site. Together, these results support a single-introduction event. We consider implications of these findings for the genetic paradox of invasion and discuss priorities for future research in *D. similis*, a promising model system for invasion biology.

## Introduction

Human-mediated biological introductions are a ubiquitous part of changing global ecosystems with negative consequences for local flora and fauna (Carlton and Geller 1993, Wonham et al. 2001, Simberloff 2013, Capinha et al. 2015). Biological introductions involving plant-feeding (phytophagous) insects are particularly common and can cause widespread damage to local crops and plants (Schulz et al. 2020, Lesieur et al. 2019). This destruction is exacerbated when insects adapt to native plant hosts, which can lead to rapid range expansion and added complications for controlling the invasion (Kennedy and Storer 2000). For this reason, understanding how and why host shifts occur in invasive species is of considerable applied importance (Clavero and Garcia-Berthou 2005, Oerke 2005). From a basic research perspective, invasive species are unintentional “evolutionary experiments” that enable us to better understand the genetic and evolutionary mechanisms underlying rapid host adaptation (North et al. 2021, Forister et al. 2012, Futuyma and Moreno 1988, Lee 2002, Vertacnik and Linnen 2017, Prentis et al. 2008). Despite considerable research effort (Bock et al. 2015, North et al. 2021) many questions regarding evolution in invasive populations remain unresolved.

One unresolved question in invasion biology asks: Given the profound reduction in genetic variation that accompanies many species introductions, how do invading populations have sufficient raw genetic material to adapt to novel environments (Allendorf & Lundquist 2003, Frankham 2005)? In recent years, the prevalence of this so-called ‘invasion paradox’ has been challenged (Estoup et al. 2016, Dlugosch and Parker 2008), with several studies failing to find evidence of reduced diversity in recently introduced populations (Roman and Darling 2007, Facon et al. 2008, Kolbe et al. 2004—but see Kanuch et al. 2021). A common mechanism for maintaining high levels of genetic variation in invasive populations is admixture between multiple genetically distinct invading populations (Bock et al. 2015, Dlugosch and Parker 2008, Prentis et al. 2008, Rius and Darling 2014). Most evidence used to support a multiple-invasion scenario is the existence of multiple genetically distinct groups within the introduced range, often inferred via patterns of population structure (Jaspers et al. 2021, Fitzpatrick et al. 2012, Sherpa et al. 2019). However, population structure can be missed when too few individuals or genetic markers are sampled (Sherpa et al. 2018, Lewald 2021). To reconstruct the demographic history of invasions and identify recent targets of selection, genome-scale data (e.g., whole-genome re-sequencing (WGS) data, are ideal (North et al. 2021). Analysis of WGS data is greatly facilitated by high quality reference genomes; however, genomic resources are still lacking for many insects (Hotaling et al. 2021, North et al. 2021, Ekblom and Galindo 2011).

Here, we develop genomic resources and describe population structure for invasive populations of *Diprion similis* (Hymenoptera: Diprionidae), a potentially powerful model for invasion population genomics. *D. similis* is an ideal organism for studying adaptation following invasion: it has undergone a pronounced host shift in the invasive range, and the introduction and spread of this species is well documented in literature and museum collections (Britton 1915, Britton 1916, Middleton 1923). *D. similis* was first recorded in Connecticut, 1914 and rapidly spread across eastern North. In its native Eurasian range, *D. similis* is found primarily on the thick and resinous needled Scots pine, *Pinus sylvestris*. In North America, however, *D. similis* is mostly found on a native, thin-needled pine, eastern white pine (*Pinus strobus*) (Lyons 2014, Coppel et al. 1974). This host association is common enough that North American *D. similis* is now casually referred to as the “introduced white-pine sawfly” by the research community. The shift from a thick-needled Eurasian host to a strikingly thin-needled North American host is somewhat surprising because thin needles represent a substantial ecological barrier to successful reproduction in many diprionids. This is because sawfly females must embed their eggs within needles without cutting too deeply, or eggs will fail to develop (Figure 1B; Knerer and Atwood 1973, McCullough and Wagner 1993, Codella and Raffa 2002, Bendall et al. 2017). Indeed, although white pine is widespread and abundant in eastern North America, only a single native diprionid, *Neodiprion pinetum*, uses this host regularly (Linnen and Farrell 2010). *N. pinetum* has also evolved several adaptations for laying eggs in thin needles, including smaller eggs, smaller ovipositors, and a unique egg-laying pattern (Bendall et al. 2017, Bendall et al. 2022, Glover et al. in prep).

**Figure 1:**
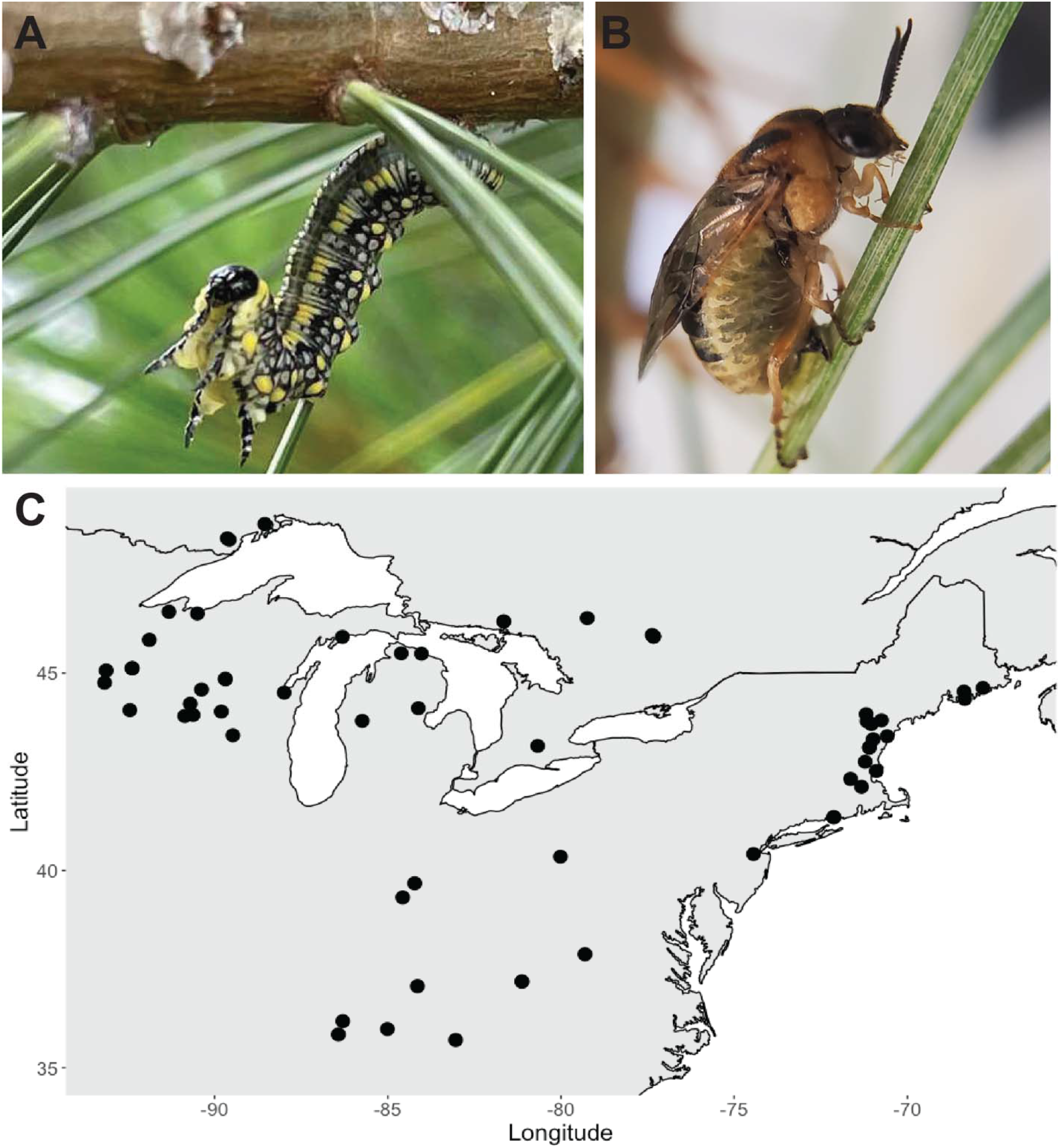
**A)** Photo of late-instar *Diprion similis* larva on white pine. Photo by Jane Dostart. **B)** Photo of *D. similis* adult female ovipositing eggs on white pine (*Pinus strobus*). **C)** Sampling locations of *Diprion similis* in eastern North America (United States and Canada) used for WGS.

As a first step to understanding how invasive *D. similis* populations were able to adapt to a novel and particularly challenging host, we describe patterns of population structure in the invasive range. To do so, we assemble a high-quality reference genome for this species, to which we map low-coverage whole-genome sequence data from 64 diploid *D. similis* individuals sampled across eastern North America. With these data, we examined three spatial patterns of genetic variation to evaluate support for single-invasion vs. multiple-invasion scenarios. First, a single introduction is expected to yield a single genetic cluster within the introduced range, while multiple invasions would be detectable as genetically distinct groups (van Boheeman et al. 2017, Sherpa et al. 2019). Second, in a single-introduction scenario—and assuming sufficient time for invasive populations to reach drift-migration equilibrium—genetic distance between individuals in the introduced range is expected to follow a mostly continuous pattern of isolation-by-distance, whereas discontinuities in the spatial distribution of genetic variation would be indicative of multiple recent invasions (Sherpa et al. 2018). Third, under a single-invasion scenario, genetic diversity should decrease from the point of introduction due to repeated founder effects and consistently smaller population sizes on the edge of range expansion; by contrast, a multiple-invasion scenario with multiple points of introduction would increase diversity and disrupt spatial patterns of genetic diversity (Van Petegem et al. 2017, Biolozyt et al. 2005, Hewitt 1993). By evaluating our data considering these predictions, we conclude that sampled North American populations of *D. similis* most likely came from a single invasion. In the discussion, we consider limitations of our data and explanations for how these populations were able to adapt to novel hosts in the absence of an influx of genetic variation via admixture.

## Materials and Methods

### DNA extraction and library preparation

For assembling a reference genome, genomic DNA from *D. similis* was isolated from one haploid male eonymph that was flash frozen in liquid nitrogen. Genomic nucleic acid isolation was performed using the MagAttract HMW DNA Kit (Qiagen, Hilden Germany) precisely following the instructions of the fresh or frozen tissue protocol. After isolation, a 2.0x Solid Phase Reverse Immobilisation (SPRI) bead clean-up was performed to improve sample purity. At each step, double-stranded DNA was quantified using the dsDNA Broad Range (BR) Qubit assay and the fluorometer feature of a DS-11 Spectrophotometer and Fluorometer (DeNovix Inc, Wilmington, DE, USA). Sample purity was determined with the UV-Vis spectrometer feature on the DS-11 which reported OD 260/230 and 260/280 ratios. Isolated genomic DNA was sheared to a mean size distribution of 20 kb using a Diagenode Megaruptor 2 (Denville, New Jersey, USA) and fragment size was confirmed using a Fragment Analyzer (Agilent Technologies, Santa Clara, California, USA) using the High Sensitivity (HS) Large Fragment kit. The sheared DNA was used for PacBio SMRTBell library preparation using the SMRTbell Express Template Prep Kit 2.0 (Pacific Biosciences, Menlo Park, California, USA) according to the manufacturer protocol. The finished library was sequenced at the USDA-ARS Genomics and Bioinformatics Research Unit in Stoneville, Mississippi, USA, where the polymerase binding reaction was performed and sequencing was carried out on one Pacific Biosciences 8M SMRT Cell on a Sequel IIe system (Pacific Biosciences, Menlo Park, California, USA) using a 30-hour movie collection time after a 2-hour pre-extension. Following sequence collection, consensus sequences from the raw PacBio Sequel IIe subreads was called using the SMRTLink v8.0 software (Pacific Biosciences, Menlo Park, California, USA).

To complement the PacBio HiFi sequencing, enriched chromosome conformation capture (HiC) sequencing was performed using another *D. similis* eonymph male sample from the same clutch of progeny from a single unmated *D. similis* female (*D. similis* is arrhenotokous, and virgin females produce haploid, male-only families). The Arima HiC kit (Arima Genomics, San Diego, California, USA) was used to perform the proximity ligation after initial crosslinking using the Arima HiC low input protocol. Proximity-ligated DNA was sheared using a Bioruptor (Diagenode, Dennville, New Jersey, USA), and appropriately sized fragments (200-600 bp) were selected as the input for the Illumina library preparation using the Swift Accel NGS 2S Plus kit (Integrated DNA Technologies, Coralville, Iowa, USA). The final library was sequenced on a fraction of a lane of NovaSeq 6000 with 150 bp paired-end sequencing. Adapter trimming was performed using BaseSpace software (Illumina, San Diego, California, USA).

### Genome assembly

Adapter-contaminated HiFi reads were filtered from the circular consensus sequencing (CCS) dataset using HiFiAdapterFilt v2.0 (Sim et al, 2022). Filtered CCS reads were assembled into a contig assembly using HiFiASM v0.16.1-r375 (Cheng et al. 2021) with no modifications to the default parameters. The default output of HiFiASM is a contig assembly displayed in graphical fragment assembly (gfa) format which was converted to fasta format using any2fasta (Seeman, 2018). Due to the extremely high contiguity of the contig assembly, the assembly was manually scaffolded using HiC data, employing the Phase Genomics HiC functions (Kronenberg and Sullivan, 2018, phasegenomics.github.io, https://github.com/phasegenomics/juicebox_scripts) (Phase Genomics, Seattle, Washington, USA) and Juicebox assembly tools (Dudchenko et al. 2017). The HiC reads were mapped to the contig assembly using bwa mem and purged of PCR duplicate artifacts using samblaster. The resulting bam file was converted into a hic formatted file using a combination of Matlock, which generates a links file, which is then converted to a .hic file using ‘run-assembly-visualizer.sh’ from 3d-dna. The HiC data was then used to manually join and edit contigs into chromosomes using Juicebox v1.11.08 (Durand et al. 2015) and the edits were applied to the contig assembly using juicebox_assembly_converter.py by Phase Genomics.

### Assembly quality analysis

Assembly quality was assessed using metrics for completeness in terms of gene content, artifact duplicate content, parity with estimated genome size, and taxonomic assignment of each assembled fragment. Completeness and amount of duplication were assessed by identifying presence of a benchmark of universal single-copy ortholog (BUSCO) genes from the Eukaryota, Metazoa, Arthopoda, Insecta, Endopterygota, and Hymenoptera odb10 databases through *ab initio* gene prediction of the assembly using Metaeuk v.4.a0f584d (Levy et al. 2020) for all the ortholog databases except for the Metazoa database which required annotation using Augustus v3.4.0 (Stanke et al. 2008). Annotations and designations of whether the orthologous genes were found in complete single copy, duplicated, fragmented, or missing were evaluated using BUSCO v5.2.2 (Manni et al. 2021) in ‘genome’ mode. Genome size was estimated using k-mer abundance calculated using KMC v3.2.1 (Kokot et al. 2017) and analyzed using GenomeScope v.2.0 (Ranallo-Benavidez, 2020). Genome coverage was estimated using KAT v2.4.2 (Mapleson et al. 2017) which uses k-mer abundance and spectra analysis to assess ploidy, coverage depth, and amount of duplication in the assembly relative to the raw reads. Finally, taxonomic assignment of each assembly scaffold or contig was determined by local alignment using nucleotide-nucleotide BLAST v2.5.9+, ‘blastn’ (Camacho et al. 2009) to the NCBI nt database (accessed on 2017-06-05) and Diamond v.2.0.9.147, ‘diamond blastx’ (Buchfink et al. 2021) to the UniProt Reference Proteomes database (accessed on 2020-03). Local alignment results were used to assign scaffolds and contigs to taxa using the blobtools v.2.6.1(Challis, et al. 2020) taxrule function ‘bestsumorder’ which assigns taxonomic identity first based on nucleotide BLAST hit followed by the preoteome BLAST hit. Scaffold and contig coverage was determined by mapping the raw reads back to the assembly using minimap2 v2.2-r1101 (Li, 2021). Results from the BUSCO analyses, alignments to the nucleotide and proteome databases, and read coverage were summarized using blobblurb (Sim, 2022).

### WGS Sample collection, DNA extraction, library prep, and sequencing

We extracted and sequenced DNA from 84 *Diprion similis* individuals collected across 77 sites on six different pine hosts (Table S1). Our sampling scheme was chosen to maximize the geographic and host range of available preserved samples, consistent with our overall goal of evaluating broad-scale demographic patterns across the introduced range. All samples were obtained from preserved larvae or adult females collected between 2001 and 2020 and stored in 95-100% ethanol at −20 °C until use. Individuals were dissected to avoid contamination from the midgut (in the case of larvae) or eggs (in the case of adult females) and then DNA was extracted using Qiagen DNeasy Tissue Kits (Qiagen Inc., Valencia, CA, USA). DNA concentrations were then assessed using a Qubit 2.0 fluorometer (Invitrogen, Waltham, MA, USA).

Library preparation and next-generation sequencing were performed on all 84 samples at Admera Health (South Plainfield, NJ, USA). Library prep was performed using KAPA HyperPrep (Roche, Basel, Switzerland) kits. Each sample was sequenced using 150 bp paired-end reads on an Illumina Novaseq 6000 S4 flowcell (Illumina, San Diego, CA).

### Data filtering: hard-genotype calls, contamination, and haploid males

Raw, demultiplexed reads were first processed using Trimmomatic (v0.3; Bolger et al. 2014) to trim adapter sequences from the ends of reads. Reads were then aligned to the *Diprion similis* reference genome using the end-to-end alignment option of Bowtie2 (v2.2.3, Langmead and Salzberg 2012). We then used Samtools (v0.1.19; Li et al. 2009) to exclude reads that mapped to more than one location in our reference genome.

Because downstream analyses assume diploidy for all samples and diprionid sawflies— like all hymenopterans—have haplodiploid sex determination, we removed putative haploid males from our dataset. However, most of our samples were preserved larvae, which we cannot sex reliably, and pine-sawfly males can occasionally be diploid (Harper et al. 2016; Bagley et al. 2017). We therefore used heterozygosity to infer ploidy. To do so, we obtained hard-genotype calls using mpileup from bcftools (v1.19, Li et al. 2011) and the -het option in vcftools (v1.19 Danacek et al. 2011). While processing these data, we found evidence of small amounts of contamination in samples—indicated by skewed allele counts unlikely to be the result of true heterozygosity. To address this, sites with skewed allele counts were flagged in each individual as missing data for downstream filtering. We then removed any site where more than half the individuals had missing data, retaining only sites with 5x individual depth of coverage and a minimum base Phred score of 20. After filtering for contamination, we examined patterns of heterozygosity across individuals, we excluded 20 individuals with <0.02 heterozygosity as putative haploid males, for a final dataset of 64 diploid individuals (Table S2).

### ANGSD genotype-likelihood estimation

To account for genotype uncertainty in downstream analyses—a recommended strategy for analyzing WGS datasets with coverage as low as 1x (Lou et al. 2020) —we used ANGSD (v0.933, Korneliussen et al. 2014). This program produces genotype-likelihood estimates in lieu “hard” single-nucleotide polymorphism (SNP) calls, and these genotype likelihoods were used for most downstream analyses (but see below). For our dataset, genotype likelihoods were estimated after keeping sites where >50% of samples passed filters for minimum mapping quality and base quality of 20, minor allele frequency > 0.05, minimum coverage depth of 1x, and maximum coverage depth of 100x (to remove repetitive loci). We then pruned using linkage disequilibrium calculated from genotype likelihoods using ngsLD (v1.1.1, Fox et al. 2019), which estimates linkage disequilibrium across the genome to produce a list of unlinked SNPs. With this list, we used ANGSD to estimate genotype likelihoods only for unlinked SNPs for use in downstream analyses. However, two analyses—isolation by distance and heterozygosity (see below)—required “hard” SNP calls instead of likelihoods. To facilitate this, ANGSD was re-run with the -dobcf and -dogeno options to produce a bcf file with “hard” SNPs at the same sites as the genotype-likelihood approach. This approach kept all the same sites as used in the genotype-likelihood approach, as these sites were already filtered.

### Evaluation of discrete population structure: PCA and NgsAdmix

Population structure within the introduced range of *D. similis* was inferred by two individual-based approaches that use genotype likelihoods and are implemented as extensions of the ANGSD platform. First, we used the program PCAngsd (v1.03, Meisner and Albrechtsen 2018) to estimate the genetic covariance matrix and perform a principal component analysis (PCA) on low-coverage genotype-likelihood data. This approach allows visualization and analysis of genetic clustering via admixture estimation from principal axes of genetic variation. Based on patterns revealed in this and other structure analyses (see below and *Results*), we also used the *lm* function of base R (version 4.2.0 R Core Team 2022) to model the first principal component eigenvector (PC1) as a function of geography (longitude of sample location). To infer the number of populations (*K*) based on the PCA analysis, we chose the value of *K* that was one fewer than the number of eigenvalues that pass Velicier’s minimum average partial (MAP) test, following (Shriner 2011). To explore an alternative clustering solution, we also used the -admix command and the first 10 eigenvectors of the PCA to estimate admixture proportions for each individual for *K*=2.

For our second approach to evaluating discrete population structure, we used NgsAdmix (v33), which calculates individual admixture proportions from low-coverage NGS data by accounting for uncertainty present in genotype likelihoods (Skotte et al. 2013). Because we have no *a priori* prediction for *K*, we ran NgsAdmix with a range of *K* values from 1 to 7, with 10 independent runs for each value of *K*. We then computed the average resulting likelihoods for each *K* value to evaluate the most likely optimal *K* value. As was done for the PCA-based approach, we also examined admixture proportions for the second-best clustering solution, *K*=2.

### Evidence of continuous population structure (isolation-by-distance)

To determine whether there is evidence of isolation-by-distance (IBD) in the introduced range, we first generated geographic and genetic matrices using SPAGEDI (v1.5b; Hardy & Vekemans 2002). Because our individuals were not sampled from discrete populations, we calculated pairwise genetic distances Rousset’s â which is analogous to the *F*_ST_/(1 – *F*_ST_) ratio (Rousset 2000). Briefly, for a pair of individuals i and j, Rousset’s distance â is given by a_ij_ = (Q_w_-Q_ij_)/(1 –Q_w_), where Q_ij_ is the probability of identity by state of gene copies between individuals and Q_w_ is the probability of identity within individuals (estimated from all pairs of individuals in the sample). We calculated pairwise Rousset’s â (Rousset 2000) using a set of 10,000 loci randomly downsampled from our “hard” SNP call data (see above) to comply with the maximum number of loci allowed by SPAGEDI 1.5 This program also calculated a pairwise linear geographic distance matrix between latitude and longitude coordinates of individuals. To test for IBD, we used the genetic and geographic distance matrices and the mantel.randtest() function from the ade4 package of R (v1.7, Dray and Dufour 2007) to perform a Mantel test (Mantel 1967) with 10,000 permutations.

### Spatial patterns of heterozygosity

To investigate how genetic diversity changes as a function of distance from the hypothesized location of introduction, we used individual heterozygosity estimates using the vcftools -het option on the “hard” genotype dataset (see above) as our measure of genetic diversity. For spatial distance from the origin of the invasive population, we used New Haven, Connecticut, United States (latitude: −73, longitude: 41.28) as the introduction site, in accordance with museum records of the species invasion (Britton 1915). We then calculated distance in kilometers from New Haven to each collection site using the geodist package in R (v0.0.7, Padgham et al. 2021). To evaluate the relationship between heterozygosity and distance from CT, we used the *lm* function of R to fit a linear model to the data.

## Results

### Diprion similis de novo *genome assembly*

The *D. similis* iyDipSimi1.1 (NCBI project: <u>JAJNQI01</u>) haploid assembly was sequenced to 100x coverage, producing an assembly size of 270.225 MB in 14 haploid chromosomes (consistent with published karyotype descriptions, Rousselet et al. 1998), 81 scaffolds, and 93 total contigs (see Table S3). The final genome size was slightly larger than the GenomeScope estimate based on k-mer abundance (Figure S1), though the larger than expected assembly was unlikely due to the inclusion of duplicate fragments (Figure S2), but rather short unplaced heterochromatic regions with an elevated GC content (Figure S3). In the final chromosome-length assembly, the smallest scaffold necessary to make up 50% of the genome (N50) was 19.014 MB, and size of the smallest scaffold necessary to make up 90% of the genome (N90) was 11.122MB (Figure S4). Completeness in terms of BUSCO annotation ranged from 91.5% of the Hymenoptera v.10 orthologs to 95.6% of the Arthropod v.10 orthologs. Of the Hymenoptera BUSCOs, all complete genes found in single copy (n = 5457 genes) or duplicated (n = 25) were in assembled chromosomes with none found in unplaced contigs. Analysis of local alignments to the NCBI nucleotide and UniProt Proteomes databases revealed no fragments from non-*D. similis* species in the assembly (Figure S3, Table S4).

### WGS sequencing, haploid filtering, and genotype-likelihood estimation

After sequencing, we obtained an average of 24.09 +/− 13.3 (SD) million reads per individual. An average of 22.59 +/− 12.84 of these reads mapped to the reference genome after removing duplicates and paralogs, and these reads covered an average of 94.3% of the reference genome. Following filtering for contamination, site depth, and base quality, we identified 22 putative haploid samples with markedly low (<0.02, Figure S5) heterozygosity that were removed from downstream analyses (Table S2). For the remaining 64 samples we filtered sites for mapping and base quality, minor allele frequency, minimum coverage of 1x, and maximum coverage of 100x, resulting in genotype likelihoods for 728,627 variable sites. After pruning based on linkage disequilibrium, we retained 352,385 genotype likelihoods for unlinked SNPs for downstream analysis. An analogous dataset with “hard” genotyped SNPs at the same sites was also used for IBD and heterozygosity analyses.

### Discrete population structure, isolation-by-distance, and heterozygosity

Analyses using PCAangsd and NgsAdmix both selected *K*=1 as the optimal number of genetic clusters in this dataset. For the PCA-based approach, *K*=1 was supported by the observation that only two eigenvectors passed the MAP test (Shriner 2011). Visualization of these two eigenvectors (principal components) indicates that much of the variation is explained by PC1 (10.5% of overall variance, Figure 2A), which correlates strongly with geography (linear model: *F* = 69.2, *P* < 0.001, *R*^2^ = 0.527, Figure 2B). PC2 appears to explain additional variation among individuals sampled from the eastern portion of the range, closer to the presumed invasion site, but no strong geographic patterns emerge from further dissecting this axis (Figure 1C). For the NgsAdmix analysis, *K*=1 was supported by likelihood values from 10 replicate runs, which matched our results from PCAngsd (Figure S6).

**Figure 2:**
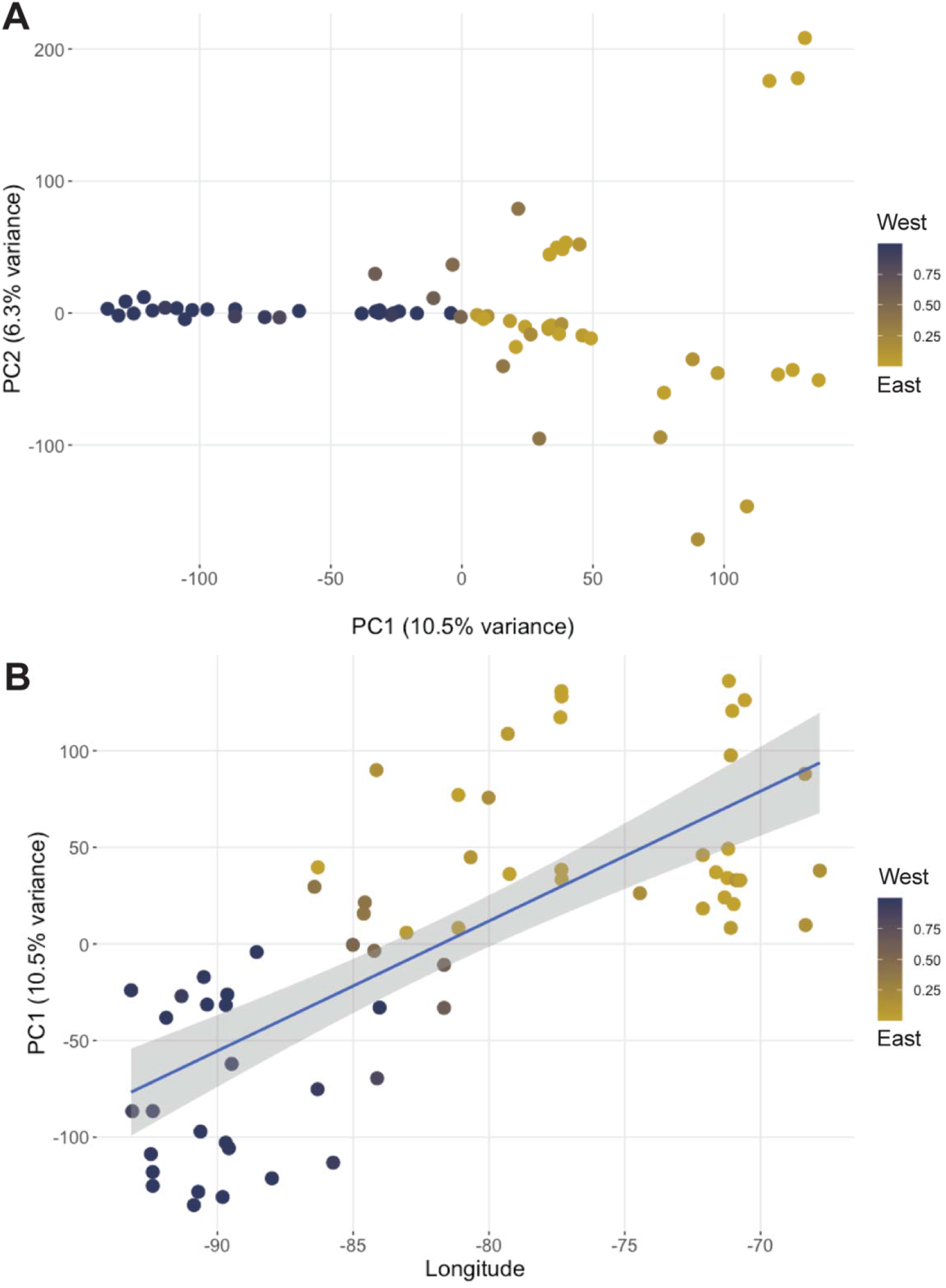
Results of PCAgsd analysis of population structure. **A)** Principal component axes of genetic variation from samples, as calculated by PCAngsd. The color shading of individual points in both panels shows admixture proportion as determined by PCAngsd for K=2 and analogous to the admixture proportions shown in Figure S7 Note that K=1 is the optimal clustering solution, admixture proportions for K=2 are shown to highlight the lack of a discrete break corresponding to two clusters. Panel A shows the component axes with highest contribution of overall variance, with PC1 inverted to align better with geographic orientation. **B** PC1 as a function of longitude of origin for each sample, and these variables are significantly correlated (linear model: *F* = 69.2, *P* < 0.001, *R*^2^ = 0.5274).

To further evaluate the potential for substructure in the data, we also evaluated population assignments (admixture proportions) for the next best clustering solution, *K*=2. Both PCAngsd and NgsAdmix produced very similar admixture proportions, with a continuous transition in admixture proportions between eastern and western groups (Figure S7). The lack of a discrete break between the two groups for the *K*=2 clustering solution (Figure S7) is consistent with a pattern of isolation-by-distance in a single continuous population (Miermans 2012).

To evaluate evidence for isolation-by-distance in the introduced range, we used Mantel tests, which revealed that there was significant positive relationship between genetic distance and geographic distance (*R*^2^ = 0.562, *P* < 0.0001, Figure 3). Plotting pairwise genetic and geographic distances also revealed some discontinuities in the IBD relationship, with two somewhat distinct clusters of points (Figure 3A). The smaller cluster of points near 0 genetic distance and < 500 km from each other represents the enriched sampling in the north-west edge of *D. similis*’ range. The discontinuity seen between this and the larger cloud of points in the IBD figure might therefore be explained by gaps in sampling; increasing sampling east and south of these locations could bridge this gap.

**Figure 3:**
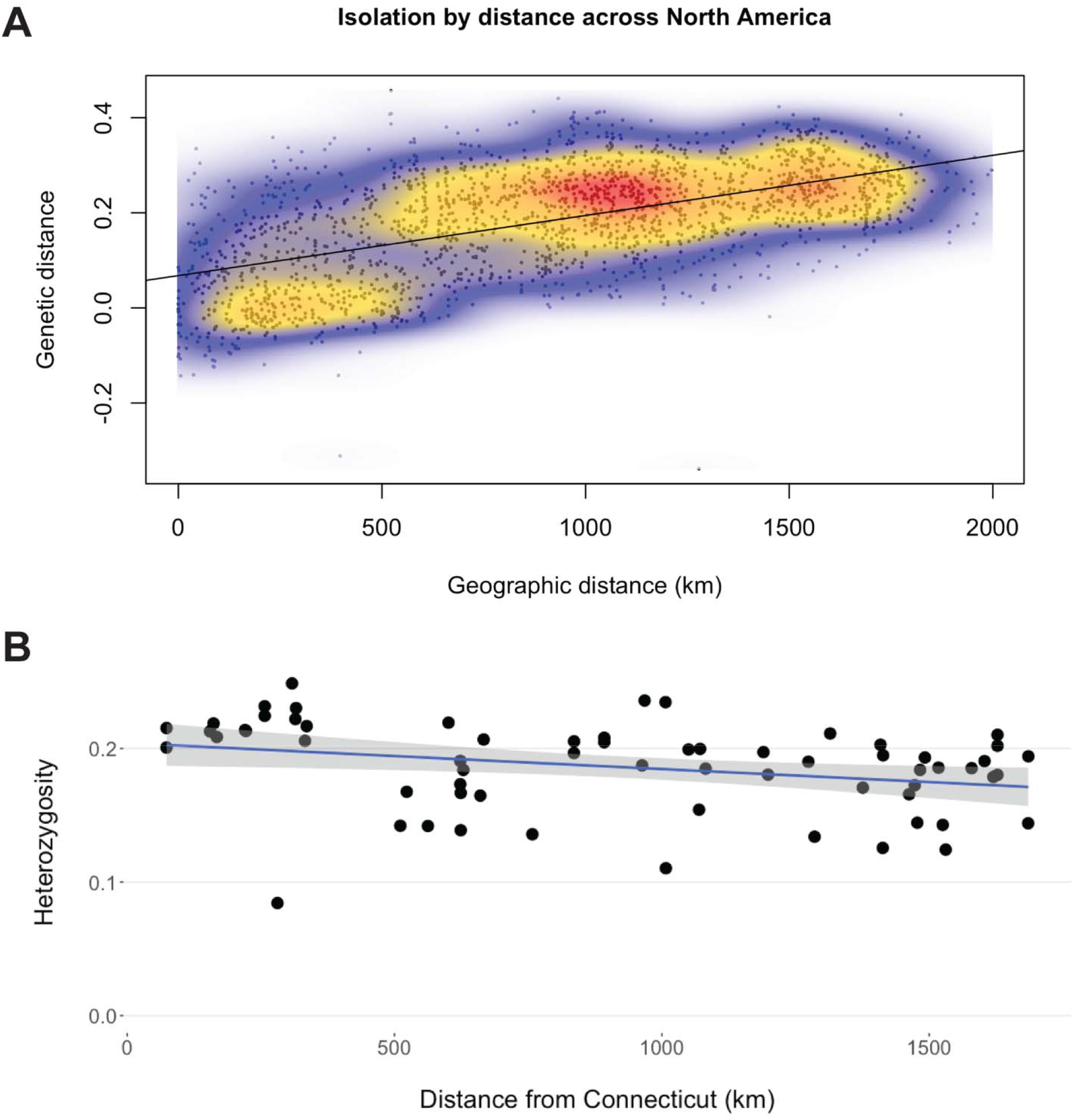
Spatial patterns of genetic variation in North American *D. similis*. **A)** Heatmap for isolation by distance (IBD) between all samples in the introduced range. Axes measure pairwise geographic and genetic distances in km and Rousset’s â respectively. Localized density between more-related individuals is indicated by warmer colors on the plot. Mantel tests indicate significant IBD range-wide (*R*^2^ = 0.562, *P* < 0.0001). These results indicate continuous isolation by distance. **B)** Spatial pattern to genetic diversity from the assumed original point of introduction in New Haven, CT. The Y-axis uses heterozygosity as a measure of genetic diversity for each individual sample. The correlation between these variables is significant (linear model: *F* = −2.47, *P* = 0.0162, *R^2^* = 0.075).

Finally, across the 64 diploid females, average observed heterozygosity was 0.186 (Figure S5). We found that heterozygosity was significantly correlated with geographic distance from the presumed location of first introduction in New Haven, CT, USA (*F* = −2.47, *P* = 0.0162, *R*^2^ = 0.075), with individuals further from this point showing reduced diversity (Figure 3B).

## Discussion

Analyses of population structure in successful biological invasions are essential for understanding the demographic and evolutionary processes behind these “natural experiments” (Lee 2002, Sakai et al. 2001, Yoshida et al. 2007). But accurate analysis of population structure may require genomic datasets consisting of many unliked markers (Sherpa et al. 2018, Lewald et al. 2021). Moreover, if detection of locations in the genome that have undergone recent positive selection is a goal, whole-genome data are better suited to the task than reduced-representation data (reviewed North et al. 2021). As a first step to developing *Diprion similis* as a model system for invasion biology, we assembled a high-quality reference genome and generated low-coverage WGS data for 64 diploid (female) individuals sampled across the introduced range of *D. similis* in North America. We found strong evidence for a single North American population of this species, with a pattern of isolation-by-distance in the introduced range. Here, we discuss both the limitations and implications of our data, while also highlighting future directions that leverage *D. similis* as a model for investigating genomics of adaptation following biological introduction.

### Spatial patterns of genetic variation support a single-invasion scenario

Our main purpose for describing population structure in North American *D. similis* was to distinguish between single-invasion and multiple-invasion scenarios, an essential first step to understanding how invasive populations adapted to a novel pine, *Pinus strobus*. We evaluated three predictions for a single-invasion scenario. First, a single wave of invasion should yield population structure with a single genetic cluster in the introduced range, while subsequent invasions are readily identified as separate genetic groups (van Boheeman et al. 2017). Our analyses identified *K*=1—a single genetic cluster—as the mostly likely number of populations within the introduced range of *D. similis*. Further supporting a single-invasion scenario, population assignments for the next-best clustering solution (*K*=2) produced a continuous gradient of population membership rather than a discrete break between two isolated populations (Figure 2A). Second, genetic relationships between individuals in the introduced range are expected to follow a continuous isolation-by-distance (IBD) pattern only if a single invasion occurred (Sherpa et al. 2018). This predicted pattern of IBD is evident in North American *D. similis* (Figure 3A), with most admixture and gene flow occurring between spatially contiguous areas throughout the range. This indicates that in the hundred years following a single introduction, the introduced meta-population is at or near drift-migration equilibrium (Sherpa et al. 2018). Third, in a single introduction scenario, genetic diversity is expected to decrease as the invasion spreads away from the original point of introduction, due to small population sizes and genetic drift at the edges of range expansions (Van Petegem et. al 2017, Biolozyt et al. 2005, Hewitt 1993). Consistent with this, there is a subtle—but significant—decline in heterozygosity with distance from the assumed point of introduction in New Haven, CT (Figure 3B).

While our data strongly support a single introduction scenario, several limitations of our dataset should be considered. One limitation of our population structure analyses is that we have not yet sampled populations in the ancestral range, making it impossible to evaluate potential source populations for the North American *D. similis* invasion. Another limitation is that there are several small, but potentially meaningful sampling gaps across the introduced range of *D. similis* (Figure 1C). Thus, we cannot rule out the possibility that there are genetically distinct populations that we did not sample. However, apart from not having samples from very recent appearances in the northwestern USA (Looney et al. 2016), our samples span most of the North American range of *D. similis*, and our largest sampling gaps were primarily located in the middle of this range. This type of sampling gap—which would be expected to cause artificial discontinuities in allele frequencies—should bias our results to detecting more populations, not fewer (Meirmans 2012). Moreover, any unsampled populations should still be detectable via admixture with nearby populations and via disruption of IBD and heterozygosity patterns, which we do not observe apart from a minor discontinuity in our IBD plot (Figure 3A). Thus, even with some sampling gaps, our data strongly support the historical description of this invasion (Zappe 1917, Middleton 1923, Codella et al. 1991): *D. similis* was first introduced somewhere near New Haven, CT approximately 110 years ago, after which it rapidly spread over a substantial portion of eastern North America.

### Why are single-introduction scenarios rare?

Our main conclusion that the highly successful *D. similis* invasion likely stems from a single introduction event contrasts with a large body of literature demonstrating that multiple-invasion scenarios are common, and in some cases necessary for successful colonization (Blair et al. 2012, Rius et al 2012, Rosenthal et al. 2008, Kolbe et al. 2007, reviewed Rius and Darling 2014). However, most studies evaluating population genomics of invasive species are in systems with much more recent introduction events (<50 years) than the ones investigated here (>100 years), and there is some evidence that age of the invasive population may influence patterns of population structure. For example, with a history of multiple invasions of different ages in different areas, the Asian tiger mosquito *Aedes albopictus* offers insights on the relationship between invasion age and population structure. Although *A. albopictus* colonized Pacific and Indian Ocean islands more than a century ago, it was not recorded in Europe until much more recently—Albania in 1979 and Italy in 1990—where it has since expanded across the continent (Scholte and Schaffner 2007, Benedict et al. 2007). Recent work demonstrates that populations from an older invasion on Reunion Island have a continuous IBD pattern, indicative of a single introduction (Sherpa et al. 2019). In contrast, more recently established European populations have a discontinuous pattern of population structure, indicative of multiple, more recent introductions (Sherpa et al. 2018, Sherpa et al. 2019, Schmidt et al. 2020). One possible explanation for differences in population structure between old vs. young invasions is that increased trade globalization makes multiple-invasion scenarios more likely in recent years than they were in the past (Seebens et al. 2015, Sardain et al. 2019).

A non-mutually exclusive hypothesis for explaining different patterns of population structure in old vs. young invasions is that evidence of multiple introductions may get erased over time. For example, an older invasive population could have had multiple introductions, but with whole-sale extinction of early invasive populations and replacement by subsequent, more successful introductions. From a genetic variation perspective, however, extinction and replacement via a new introduction with limited admixture is essentially equivalent to a single-invasion scenario. Alternatively, if there has been enough time and sufficient gene flow between invading populations, secondary invasions may not be detectable from samples of modern invasive populations, although samples from source populations—if available—may provide some evidence of historical admixture. Overall, multiple-invasion scenarios may be much harder to detect in older invasions (e.g., >100 years) due to the possibility that introduced populations or source populations have since gone extinct. Fortunately, recent advances in museum genomics (Parejo et al. 2020, reviewed Raxworthy and Smith 2021) may make it possible to resurrect these lost populations. With good representation in natural history literature and museum collections— and now a reference genome for mapping sequencing reads from degraded samples—*Diprion similis* provides an excellent test case for using museum genomics to evaluate cryptic multiple-invasion scenarios.

### Adaptation in single-invasion scenarios

If *Diprion similis* arrived in a single introduction event as the data here suggest, it remains unclear how this species had sufficient genetic variation to shift to white pine (*Pinus strobus*) in North America. According to the genetic paradox of invasions, a single invasive wave is expected to considerably reduce genetic diversity (Allendorf and Lundquist 2003, Frankham 2005), thereby limiting the raw genetic material available for adapting to novel hosts. Nevertheless, there are several potential mitigating factors that may explain how invasive populations can adapt to novel environments despite limited genetic variation (Estoup et al. 2016). Here, we consider several non-mutually exclusive explanations that may apply to *D. similis*. The most plausible of these may be that the introduced range provides no adaptive challenge to the invading species. While *Neodiprion* pine sawfly species that do not ordinarily oviposit on white pine experience increased egg (Bendall et al. 2017) and larval (CRL, personal observation) mortality, the same may not be the case for *D. similis*. For example, unlike *Neodiprion* females, *D. similis* females use resin and pulp from the pine to cover the exposed egg (Zappe 1917, Bittner et al. 2019, Bendall et al. 2017). It is therefore possible that this difference in oviposition behavior is a pre-adaptation for white pine use in *D. similis*. One way to test this hypothesis would be to measure egg-laying success in European populations of *D. similis* that have never been exposed to *P. strobus*.

Another mechanism that could account for a rapid host shift despite presumably limited genetic variation is if novel environments in the invasive range releases phenotypic plasticity for host use (Lande 2015, Funk 2008, Zenni et al. 2014). For example, *Pinus sylvestris* is considered the primary host of *D. similis* in its native range, but it has been documented attacking a variety of other pines throughout its range (Codella et al. 1991). Across this range, *P. sylvestris* is the dominant and often sole pine species available for attack, however some locations—particularly in mountainous regions of Europe—having much greater pine diversity (Jin et al. 2021). It is therefore possible that *D. similis* populations subject to environments with greater diversity in pine hosts have increased plasticity in host-use phenotypes. Therefore, if the North American population came from a *D. similis* population with more generalized pine use, pre-existing plasticity for host acceptance—possibly coupled with pre-adaptations for laying eggs in different types of needles—may have facilitated rapid shifts to novel hosts. Testing this hypothesis will require evaluating host preference and acceptance behaviors in potential source populations within the native range of *D. similis*. Other genetic phenomena in founder populations—such as epigenetics and purging of deleterious mutations (Estoup et al. 2016)—can also promote persistence and adaptation in the invasive range. Ultimately, pairing the genomic resources generated here with additional experimental work in native and invasive *D. similis* populations would provide useful insights into the genetic mechanisms underlying host shifts in invasive populations.

## Conclusion

Overall, our results and discussion highlight the value of taking an integrative approach to evaluating the history of invasions: while genomic data are valuable, their interpretation hinges on their ecological and historical context. Here, we lay the groundwork for establishing *Diprion similis* as a model for evaluating the demographic history and genetic underpinnings of adaptation in biological invasions. Armed with the genomic resources and spatial patterns of genetic variation presented here, future work can leverage two key advantages in this system: (1) a rich collection of historical samples and data that will provide snapshots of genetic variation in past *D. similis* populations, and (2) experimental tractability for connecting genetic variation to ecologically relevant traits and their impact on fitness. Ultimately, this work will answer pressing questions about prevalence of single invasion scenarios and consequences for adaptation to novel environments.

## Data Availability

The *de novo* assembled reference genome can be found on NCBI (BioProject <u>PRJNA784632</u>). Additional data, including individual WGS sequences for all 64 samples, genotype-likelihood .beagle file, and hard-called genotype data in .vcf format can be found on Dryad repository upon publication.

## Acknowledgements

We thank current and former Linnen lab members: Robin Bagley, John Terbot, and Ashleigh Glover for collecting and providing many of the samples used in this work. We thank Jane Dostart for permission to use her photograph of a *Diprion similis* larva as well as excellent care of insects. We thank Dr. David Smith of the Smithsonian National Museum of Natural History for helpful comments on availability and richness of museum collections for this species. The authors thank the members of the USDA-ARS Ag100Pest Team for sequencing and analysis support. All opinions expressed in this paper are the author’s and do not necessarily reflect the policies and views of USDA. Mention of trade names or commercial products in this publication is solely for the purpose of providing specific information and does not imply recommendation or endorsement by the U.S. Department of Agriculture.

## References

Allendorf, F. W., & Lundquist, L. L. (2003). Introduction: Population biology, evolution, and control of Invasive Species. Conservation Biology, 17(1), 24–30. https://doi.org/10.1046/j.1523-1739.2003.02365.x

Bagley, R. K., Sousa, V. C., Niemiller, M. L., & Linnen, C. R. (2017). History, geography and host use shape genomewide patterns of genetic variation in the redheaded pine sawfly (Neodiprion lecontei). Molecular Ecology, 26(4), 1022–1044. https://doi.org/10.1111/mec.13972

Bendall, E. E., Bagley, R. K., Sousa, V. C., & Linnen, C. R. (2022). Faster-haplodiploid evolution under divergence-with-gene-flow: Simulations and empirical data from pine-feeding hymenopterans. Molecular Ecology, 31(8), 2348–2366. https://doi.org/10.1111/mec.16410

Bendall, E. E., Vertacnik, K. L., & Linnen, C. R. (2017). Oviposition traits generate extrinsic postzygotic isolation between two pine sawfly species. BMC Evolutionary Biology, 17(1), 1–15. https://doi.org/10.1186/s12862-017-0872-8

Benedict, M., Levine, R., Hawley, W., & Lounibos, L. (2007). Spread of the tiger: Global risk of invasion by the mosquito Aedes albopictus. Vector-Borne and Zoonotic Diseases, 7(1), 76–85.

Bialozyt, R., Ziegenhagen, B., & Petit, R. J. (2006). Contrasting effects of long distance seed dispersal on genetic diversity during range expansion. Journal of Evolutionary Biology, 19(1), 12–20. https://doi.org/10.1111/j.1420-9101.2005.00995.x

Bittner, N., Hundacker, J., Achotegui-Castells, A., Anderbrant, O., & Hilker, M. (2019). Defense of Scots pine against sawfly eggs (Diprion pini) is primed by exposure to sawfly sex pheromones. Proceedings of the National Academy of Sciences of the United States of America, 116(49), 24668–24675. https://doi.org/10.1073/pnas.1910991116

Blair, A. C., Blumenthal, D., & Hufbauer, R. A. (2012). Hybridization and invasion: An experimental test with diffuse knapweed (Centaurea diffusa Lam.). Evolutionary Applications, 5(1), 17–28. https://doi.org/10.1111/j.1752-4571.2011.00203.x

Bock, D. G., Caseys, C., Cousens, R. D., Hahn, M. A., Heredia, S. M., Hübner, S., Turner, K. G., Whitney, K. D., & Rieseberg, L. H. (2015). What we still don’t know about invasion genetics. Molecular Ecology, 24(9), 2277–2297. https://doi.org/10.1111/mec.13032

Britton, W. E. (1916). Further Notes on Diprion Simile Hartig. Journal of Economic Entomology, 9(2), 281–282. https://doi.org/10.1093/jee/9.2.281

Britton, W. E. (1915). A Destructive Pine Sawfly Introduced from Europe. Journal of Economic Entomology, 8(3), 379–382. https://doi.org/10.1093/jee/8.3.379

Britton WE; Zappe MP, 1918. The imported pine sawfly; Diprion (Lophyrus) simile Hartig. Conn. Agr. Exp. Sta. Bull. 203:273–290.

Buchfink, B., Reuter, K., & Drost, H. G. (2021). Sensitive protein alignments at tree-of-life scale using DIAMOND. Nature Methods, 18(4), 366–368. https://doi.org/10.1038/s41592-021-01101-x

Camacho, C., Coulouris, G., Avagyan, V., Ma, N., Papadopoulos, J., Bealer, K., & Madden, T. L. (2009). BLAST+: Architecture and applications. BMC Bioinformatics, 10, 1–9. https://doi.org/10.1186/1471-2105-10-421

Capinha, C., Essl, F., Seebens, H., Moser, D., & Miquel, H. (2022). The dispersal of alien species redefines biogeography in the Anthropocene Published by : American Association for the Advancement of Science Linked references are available on JSTOR for this article : The The dispersal dispersal of of alien alien species. Science, 348(6240), 1248–1251.

Carlton, J. T., & Geller, J. B. (1993). Ecological Roulette : The Global Transport of Nonindigenous Marine Organisms. Science, 261(5117), 78–82.

Challis, R., Richards, E., Rajan, J., Cochrane, G., & Blaxter, M. (2020). BlobToolKit - interactive quality assessment of genome assemblies. G3: Genes, Genomes, Genetics, 10(4), 1361–1374. https://doi.org/10.1534/g3.119.400908

Cheng, H., Concepcion, G. T., Feng, X., Zhang, H., & Li, H. (2021). Haplotype-resolved de novo assembly using phased assembly graphs with hifiasm. Nature Methods, 18(2), 170–175. https://doi.org/10.1038/s41592-020-01056-5.Haplotype-resolved

Clavero, M., & Garcia-Berthou, E. (2005). Invasive species are a leading cause of animal extinctions. Trends in Ecology and Evolution. TRENDS in Ecology & Evolution, 20(3), 110.

Codella, S., Fogal, W., & Raffa, K. (1991). The effect of host variability on growth and performance of the introduced pine sawfly, Diprion similis. Canadian Journal of Forest Research, 21, 1668–1674.

Codella, S. G., & Raffa, K. F. (2002). Desiccation of Pinus foliage induced by conifer sawfly oviposition: Effect on egg viability. Ecological Entomology, 27(5), 618–621. https://doi.org/10.1046/j.1365-2311.2002.00447.x

Coppel, H., Mertins, J., & Harris, J. (1974). The introduced pine sawfly, Diprion similis (Hartig) (Hymenoptera: Diprionidae). A review, with emphasis on studies in Wisconsin. University. University of Wisconsin Research Division, R2393.

Danecek, P., Auton, A., Abecasis, G., Albers, C. A., Banks, E., DePristo, M. A., Handsaker, R. E., Lunter, G., Marth, G. T., Sherry, S. T., McVean, G., & Durbin, R. (2011). The variant call format and VCFtools. Bioinformatics, 27(15), 2156–2158. https://doi.org/10.1093/bioinformatics/btr330

Dlugosch, K. M., & Parker, I. M. (2008). Founding events in species invasions: Genetic variation, adaptive evolution, and the role of multiple introductions. Molecular Ecology, 17(1), 431–449. https://doi.org/10.1111/j.1365-294X.2007.03538.x

Dlugosch, K. M., Anderson, S. R., Braasch, J., Cang, F. A., & Gillette, H. D. (2015). The devil is in the details: Genetic variation in introduced populations and its contributions to invasion. Molecular Ecology, 24(9), 2095–2111. https://doi.org/10.1111/mec.13183

Dray, S., & Dufour, A. B. (2007). The ade4 package: Implementing the duality diagram for ecologists. Journal of Statistical Software, 22(4), 1–20. https://doi.org/10.18637/jss.v022.i04

Dudchenko, O., Batra, S. S., Omer, A. D., Nyquist, S. K., Hoeger, M., Durand, N. C., Shamim, M. S., Machol, I., Lander, E. S., Aiden, A. P., & Aiden, E. L. (2017). De novo assembly of the Aedes aegypti genome using Hi-C yields chromosome-length scaffolds. Science, 356(6333), 92–95. https://doi.org/10.1126/science.aal3327.De

Durand, N. C., Shamim, M. S., Machol, I., Rao, S. S. P., Huntley, M. H., Lander, E. S., & Aiden, E. L. (2016). Juicer Provides a One-Click System for Analyzing Loop-Resolution Hi-C Experiments. Cell Systems, 3(1), 95–98. https://doi.org/10.1016/j.cels.2016.07.002

Ekblom, R., & Galindo, J. (2011). Applications of next generation sequencing in molecular ecology of non-model organisms. Heredity, 107(1), 1–15. https://doi.org/10.1038/hdy.2010.152

Estoup, A., Ravigné, V., Hufbauer, R., Vitalis, R., Gautier, M., & Facon, B. (2016). Is There a Genetic Paradox of Biological Invasion? Annual Review of Ecology, Evolution, and Systematics, 47, 51–72. https://doi.org/10.1146/annurev-ecolsys-121415-032116

Facon, B., Pointier, J. P., Jarne, P., Sarda, V., & David, P. (2008). High Genetic Variance in Life-History Strategies within Invasive Populations by Way of Multiple Introductions. Current Biology, 18(5), 363–367. https://doi.org/10.1016/j.cub.2008.01.063

Fitzpatrick, B. M., Fordyce, J. A., Niemiller, M. L., & Reynolds, R. G. (2012). What can DNA tell us about biological invasions? Biological Invasions, 14(2), 245–253. https://doi.org/10.1007/s10530-011-0064-1

Forister, M. L., Dyer, L. A., Singer, M. S., Stireman, J. O., & Lill, J. T. (2012). Revisiting the evolution of ecological specialization, with emphasis on insect-plant interactions. Ecology, 93(5), 981–991. https://doi.org/10.1890/11-0650.1

Fox, E. A., Wright, A. E., Fumagalli, M., & Vieira, F. G. (2019). NgsLD: Evaluating linkage disequilibrium using genotype likelihoods. Bioinformatics, 35(19), 3855–3856. https://doi.org/10.1093/bioinformatics/btz200

Frankham, R. (2005). Resolving the genetic paradox in invasive species. Heredity, 94(4), 385. https://doi.org/10.1038/sj.hdy.6800634

Funk, J. L. (2008). Differences in plasticity between invasive and native plants from a low resource environment. Journal of Ecology, 96(6), 1162–1173. https://doi.org/10.1111/j.1365-2745.2008.01435.x

Futuyma, D. J., & Moreno, G. (1988). The Evolution of Ecological Specialization Author (s): Douglas J. Futuyma and Gabriel Moreno Source : Annual Review of Ecology and Systematics, Vol. 19 (1988), pp. 207–233 Published by : Annual Reviews Stable URL: http://www.jstor.org/stable/2097. Annual Review of Ecology and Systematics, 19(1988), 207–233.

Hardy, O., & Vekemans, X. (2002). Spagedi : a Versatile Computer Program To Analyse Spatial. Molecular Ecology Notes, 2, 618–620. https://doi.org/10.1046/j.1471-8278

Harper, K. E., Bagley, R. K., Thompson, K. L., & Linnen, C. R. (2016). Complementary sex determination, inbreeding depression and inbreeding avoidance in a gregarious sawfly. Heredity, 117(5), 326–335. https://doi.org/10.1038/hdy.2016.46

Hewitt, G. M. (1999). Post-glacial re-colonization of European biota. Biological Journal of the Linnean Society, 68(1–2), 87–112. https://doi.org/10.1006/bijl.1999.0332

Hotaling, S., Sproul, J. S., Heckenhauer, J., Powell, A., Larracuente, A. M., Pauls, S. U., Kelley, J. L., & Frandsen, P. B. (2021). Long Reads Are Revolutionizing 20 Years of Insect Genome Sequencing. Genome Biology and Evolution, 13(8), 1–7. https://doi.org/10.1093/gbe/evab138

Jaspers, C., Ehrlich, M., Pujolar, J. M., Kunzel, S., Bayer, T., Limborg, M. T., Lombard, F., Browne, W. E., Stefanova, K., & Reusch, T. B. H. (2021). Invasion genomics uncover contrasting scenarios of genetic diversity in a widespread marine invader. Proceedings of the National Academy of Sciences of the United States of America, 118(51). https://doi.org/10.1073/pnas.2116211118

Jin, W. T., Gernandt, D. S., Wehenkel, C., Xia, X. M., Wei, X. X., & Wang, X. Q. (2021). Phylogenomic and ecological analyses reveal the spatiotemporal evolution of global pines. Proceedings of the National Academy of Sciences of the United States of America, 118(20). https://doi.org/10.1073/PNAS.2022302118

Kaňuch, P., Berggren, Å., & Cassel-Lundhagen, A. (2021). A clue to invasion success: genetic diversity quickly rebounds after introduction bottlenecks. Biological Invasions, 23(4), 1141–1156. https://doi.org/10.1007/s10530-020-02426-y

Kennedy, G. G., & Storer, N. P. (2000). LIFE SYSTEMS OF POLYPHAGOUS ARTHROPOD PESTS IN TEMPORALLY UNSTABLE CROPPING SYSTEMS. Annual Review of Entomology, 45, 467–493.

Knerer, A. G., & Atwood, C. E. (2018). Diprionid Sawflies : Polymorphism and Speciation. Science, 179(4078), 1090–1099.

Kokot, M., Dlugosz, M., & Deorowicz, S. (2017). KMC 3: counting and manipulating k-mer statistics. Bioinformatics (Oxford, England), 33(17), 2759–2761. https://doi.org/10.1093/bioinformatics/btx304

Kolbe, J. J., Glor, R. E., Schettino, L. R. G., Lara, A. C., Larson, A., & Losos, J. B. (2004). Genetic variation increases during a biological invasion. Nature, 431(7005), 177–181.

Korneliussen, T. S., Albrechtsen, A., & Nielsen, R. (2014). ANGSD: Analysis of Next Generation Sequencing Data. BMC Bioinformatics, 15(1), 1–13. https://doi.org/10.1186/s12859-014-0356-4

Kronenburg, Z., & Sulllivan, S. (2017). Matlock. https://github.com/phasegenomics/matlock

Lande, R. (2015). Evolution of phenotypic plasticity in colonizing species. Molecular Ecology, 24(9), 2038–2045. https://doi.org/10.1111/mec.13037

Langmead, B., & Salzberg, S. L. (2012). Fast gapped-read alignment with Bowtie 2. Nature Methods, 9(4), 357–359. https://doi.org/10.1038/nmeth.1923

Lee, C. E. (2002). Evolutionary genetics of invasive species. Trends in Ecology and Evolution, 17(8), 386–391. https://doi.org/10.1016/S0169-5347(02)02554-5

Lesieur, V., Lombaert, E., Guillemaud, T., Courtial, B., Strong, W., Roques, A., & Auger-Rozenberg, M. A. (2019). The rapid spread of Leptoglossus occidentalis in Europe: a bridgehead invasion. Journal of Pest Science, 92(1), 189–200. https://doi.org/10.1007/s10340-018-0993-x

Levy Karin, E., Mirdita, M., & Söding, J. (2020). MetaEuk-sensitive, high-throughput gene discovery, and annotation for large-scale eukaryotic metagenomics. Microbiome, 8(1), 1–15. https://doi.org/10.1186/s40168-020-00808-x

Lewald, K. M., Abrieux, A., Wilson, D. A., Lee, Y., Conner, W. R., Andreazza, F., Beers, E. H., Burrack, H. J., Daane, K. M., Diepenbrock, L., Drummond, F. A., Fanning, P. D., Gaffney, M. T., Hesler, S. P., Ioriatti, C., Isaacs, R., Little, B. A., Loeb, G. M., Miller, B., … Chiu, J. C. (2021). Population genomics of Drosophila suzukii reveal longitudinal population structure and signals of migrations in and out of the continental United States. G3: Genes, Genomes, Genetics, 11(12). https://doi.org/10.1093/g3journal/jkab343

Li, H. (2021). New strategies to improve minimap2 alignment accuracy. Bioinformatics, 37(23), 4572–4574. https://doi.org/10.1093/bioinformatics/btab705

Li, H. (2011). A statistical framework for SNP calling, mutation discovery, association mapping and population genetical parameter estimation from sequencing data. Bioinformatics, 27(21), 2987–2993. https://doi.org/10.1093/bioinformatics/btr509

Linnen, C. R., & Farrell, B. D. (2010). A test of the sympatric host race formation hypothesis in Neodiprion (Hymenoptera: Diprionidae). Proceedings of the Royal Society B: Biological Sciences, 277(1697), 3131–3138. https://doi.org/10.1098/rspb.2010.0577

Looney, C., Smith, D. R., Collman, S. J., Langor, D. W., & Peterson, M. A. (2016). Sawflies (Hymenoptera, Symphyta) newly recorded from Washington State. Journal of Hymenoptera Research, 49, 129–159.

Lyons, B. (2014). The Use of Classical Biological Control to Preserve Forests in North America: X Introduced Pine Sawfly. In USDA.

Manni, M., Berkeley, M. R., Seppey, M., Simão, F. A., & Zdobnov, E. M. (2021). BUSCO Update: Novel and Streamlined Workflows along with Broader and Deeper Phylogenetic Coverage for Scoring of Eukaryotic, Prokaryotic, and Viral Genomes. Molecular Biology and Evolution, 38(10), 4647–4654. https://doi.org/10.1093/molbev/msab199

Mantel N. (1967). The detection of disease clustering and a generalized regression approach. Cancer Research, 27(2), 209–220.

Mapleson, D., Accinelli, G. G., Kettleborough, G., Wright, J., & Clavijo, B. J. (2017). KAT: A K-mer analysis toolkit to quality control NGS datasets and genome assemblies. Bioinformatics, 33(4), 574–576. https://doi.org/10.1093/bioinformatics/btw663

McCullough, D. G., & Wagner, M. R. (1993). Defusing host defenses: ovipositional adaptations of sawflies to plant resins. Sawfly life Hist. Adapt. to woody plants, 151, 71.

Meirmans, P. G. (2012). The trouble with isolation by distance. Molecular Ecology, 21(12), 2839–2846. https://doi.org/10.1111/j.1365-294X.2012.05578.x

Meisner, J., & Albrechtsen, A. (2018). Inferring population structure and admixture proportions in low-depth NGS data. Genetics, 210(2), 719–731. https://doi.org/10.1534/genetics.118.301336

Middleton, W. (1923). The imported pine sawfly.

North, H. L., McGaughran, A., & Jiggins, C. D. (2021). Insights into invasive species from whole-genome resequencing. Molecular Ecology, 30(23), 6289–6308. https://doi.org/10.1111/mec.15999

Oerke, E. C. (2006). Crop losses to pests. Journal of Agricultural Science, 144(1), 31–43. https://doi.org/10.1017/S0021859605005708

Parejo, M., Wragg, D., Henriques, D., Charrière, J. D., & Estonba, A. (2020). Digging into the genomic past of Swiss honey bees by whole-genome sequencing museum specimens. Genome Biology and Evolution, 12(12), 2535–2551. https://doi.org/10.1093/GBE/EVAA188

Prentis, P. J., Wilson, J. R. U., Dormontt, E. E., Richardson, D. M., & Lowe, A. J. (2008). Adaptive evolution in invasive species. Trends in Plant Science, 13(6), 288–294. https://doi.org/10.1016/j.tplants.2008.03.004

R Core Team (2021). R: A language and environment for statistical computing. R Foundation for Statistical Computing, Vienna, Austria.

Ranallo-Benavidez, T. R., Jaron, K. S., & Schatz, M. C. (2020). GenomeScope 2.0 and Smudgeplot for reference-free profiling of polyploid genomes. Nature Communications, 11(1). https://doi.org/10.1038/s41467-020-14998-3

Raxworthy, C. J., & Smith, B. T. (2021). Mining museums for historical DNA: advances and challenges in museomics. Trends in Ecology and Evolution, 36(11), 1049–1060. https://doi.org/10.1016/j.tree.2021.07.009

Rius, M., & Darling, J. A. (2014). How important is intraspecific genetic admixture to the success of colonising populations? Trends in Ecology and Evolution, 29(4), 233–242. https://doi.org/10.1016/j.tree.2014.02.003

Roman, J., & Darling, J. A. (2007). Paradox lost: genetic diversity and the success of aquatic invasions. Trends in Ecology and Evolution, 22(9), 454–464. https://doi.org/10.1016/j.tree.2007.07.002

Rosenthal, D. M., Ramakrishnan, A. P., & Cruzan, M. B. (2008). Evidence for multiple sources of invasion and intraspecific hybridization in Brachypodium sylvaticum (Hudson) Beauv. in North America. Molecular Ecology, 17(21), 4657–4669. https://doi.org/10.1111/j.1365-294X.2008.03844.x

Rousselet, J., Géri, C., Hewitt, G. M., & Lemeunier, F. (1998). The chromosomes of Diprion pini and D. similis (Hymenoptera: Diprionidae): implications for karyotype evolution. Heredity, 81(5), 573–578. https://doi.org/10.1046/j.1365-2540.1998.00421.x

Sakai, A. K., Allendorf, F. W., Holt, J. S., Lodge, D. M., Molofsky, J., With, K. A., Baughman, S., Cabin, R. J., Cohen, J. E., Ellstrand, N. C., Mccauley, D. E., Neil, P. O., Parker, I. M., Thompson, J. N., & Weller, S. G. (2008). The Population Biology of Invasive Specie Source : Annual Review of Ecology and Systematics, Vol. 32, (2001), pp. 305–332 Published by : Annual Reviews Stable URL : http://www.jstor.org/stable/2678643. Ecology, 32(2001), 305–332.

Sardain, A., Sardain, E., & Leung, B. (2019). Global forecasts of shipping traffic and biological invasions to 2050. Nature Sustainability, 2(4), 274–282. https://doi.org/10.1038/s41893-019-0245-y

Schmidt, T. L., Chung, J., Honnen, A. C., Weeks, A. R., & Hoffmann, A. A. (2020). Population genomics of two invasive mosquitoes (Aedes aegypti and aedes albopictus) from the indo-pacific. PLoS Neglected Tropical Diseases, 14(7), 1–24. https://doi.org/10.1371/journal.pntd.0008463

Schmidt, T. L., Swan, T., Chung, J., Karl, S., Demok, S., Yang, Q., Field, M. A., Muzari, M. O., Ehlers, G., Brugh, M., Bellwood, R., Horne, P., Burkot, T. R., Ritchie, S., & Hoffmann, A. A. (2021). Spatial population genomics of a recent mosquito invasion. Molecular Ecology, 30(5), 1174–1189. https://doi.org/10.1111/mec.15792

Scholte, E. J., & Schaffner, F. (2007). Waiting for the tiger: Establishment and spread of the Asian tiger mosquito in Europe. Emerging Pests and Vector-Borne Diseases in Europe, 14(January 2007), 241–261.

Schulz, A. N., Mech, A. M., Allen, C. R., Ayres, M. P., Gandhi, K. J. K., Gurevitch, J., Havill, N. P., Herms, D. A., Hufbauer, R. A., Liebhold, A. M., Raffa, K. F., Raupp, M. J., Thomas, K. A., Tobin, P. C., & Marsico, T. D. (2020). The impact is in the details: Evaluating a standardized protocol and scale for determining non-native insect impact. NeoBiota, 55(August 2019), 61–83. https://doi.org/10.3897/NEOBIOTA.55.38981

Seebens, H., Essl, F., Dawson, W., Fuentes, N., Moser, D., Pergl, J., Pyšek, P., van Kleunen, M., Weber, E., Winter, M., & Blasius, B. (2015). Global trade will accelerate plant invasions in emerging economies under climate change. Global Change Biology, 21(11), 4128–4140. https://doi.org/10.1111/gcb.13021

Seeman, T. (2018). any2fasta (0.4.2). https://github.com/tseemann/any2fasta

Sherpa, S., Blum, M. G. B., Capblancq, T., Cumer, T., Rioux, D., & Després, L. (2019). Unravelling the invasion history of the Asian tiger mosquito in Europe. Molecular Ecology, 28(9), 2360–2377. https://doi.org/10.1111/mec.15071

Sherpa, S., Rioux, D., Pougnet-Lagarde, C., & Després, L. (2018). Genetic diversity and distribution differ between long-established and recently introduced populations in the invasive mosquito Aedes albopictus. Infection, Genetics and Evolution, 58(December 2017), 145–156. https://doi.org/10.1016/j.meegid.2017.12.018

Shriner, D. (2011). Investigating population stratification and admixture using eigenanalysis of dense genotypes. Heredity, 107(5), 413–420. https://doi.org/10.1038/hdy.2011.26

Sim, S. B. (2022). blobblurb. https://github.com/sheinasim/blobblurb

Sim, S. B., Corpuz, R. L., Simmonds, T. J., & Geib, S. M. (2022). HiFiAdapterFilt, a memory efficient read processing pipeline, prevents occurrence of adapter sequence in PacBio HiFi reads and their negative impacts on genome assembly. BMC Genomics, 23(1), 1–7. https://doi.org/10.1186/s12864-022-08375-1

Simberloff, D., Martin, J. L., Genovesi, P., Maris, V., Wardle, D. A., Aronson, J., Courchamp, F., Galil, B., García-Berthou, E., Pascal, M., Pyšek, P., Sousa, R., Tabacchi, E., & Vilà, M. (2013). Impacts of biological invasions: What’s what and the way forward. TRENDS in Ecology and Evolution, 28(1), 58–66. https://doi.org/10.1016/j.tree.2012.07.013

Skotte, L., Korneliussen, T. S., & Albrechtsen, A. (2013). Estimating individual admixture proportions from next generation sequencing data. Genetics, 195(3), 693–702. https://doi.org/10.1534/genetics.113.154138

Stanke, M., Diekhans, M., Baertsch, R., & Haussler, D. (2008). Using native and syntenically mapped cDNA alignments to improve de novo gene finding. Bioinformatics, 24(5), 637–644. https://doi.org/10.1093/bioinformatics/btn013

van Boheemen, L. A., Lombaert, E., Nurkowski, K. A., Gauffre, B., Rieseberg, L. H., & Hodgins, K. A. (2017). Multiple introductions, admixture and bridgehead invasion characterize the introduction history of Ambrosia artemisiifolia in Europe and Australia. Molecular Ecology, 26(20), 5421–5434. https://doi.org/10.1111/mec.14293

Van Petegem, K., Moerman, F., Dahirel, M., Fronhofer, E. A., Vandegehuchte, M. L., Van Leeuwen, T., Wybouw, N., Stoks, R., & Bonte, D. (2018). Kin competition accelerates experimental range expansion in an arthropod herbivore. Ecology Letters, 21(2), 225–234. https://doi.org/10.1111/ele.12887

Vertacnik, K. L., & Linnen, C. R. (2017). Evolutionary genetics of host shifts in herbivorous insects: insights from the age of genomics. Annals of the New York Academy of Sciences, 1389(1), 186–212. https://doi.org/10.1111/nyas.13311

Wonham, M. J., Walton, W. C., Ruiz, G. M., Frese, A. M., & Galil, B. S. (2001). Going to the source: Role of the invasion pathway in determining potential invaders. Marine Ecology Progress Series, 215, 1–12. https://doi.org/10.3354/meps215001

Yoshida, T., Goka, K., Ishihama, F., Ishihara, M., & Kudo, S. I. (2007). Biological invasion as a natural experiment of the evolutionary processes: Introduction of the special feature. Ecological Research, 22(6), 849–854. https://doi.org/10.1007/s11284-007-0435-3

Zappe, M. P. (1917). Egg-Laying Habits of Diprion Simile Hartig. Journal of Economic Entomology, 10(1), 188–190. https://doi.org/10.1093/jee/10.1.188

Zenni, R. D., Lamy, J. B., Lamarque, L. J., & Porté, A. J. (2014). Adaptive evolution and phenotypic plasticity during naturalization and spread of invasive species: Implications for tree invasion biology. Biological Invasions, 16(3), 635–644. https://doi.org/10.1007/s10530-013-0607-8

