## Supplementary data for "Whole-genome resequencing data support a single introduction of the invasive white pine sawfly, *Diprion similis*"

**Table S1: Sampling locations with latitude and longitude coordinates, as well as collection date and pine host species collected on.**

| Sample name | Date collected | Latitude | Longitude | Collection host | State |
|---|---|---|---|---|---|
| 006-01 | 7/19/01 | 42.31954 | -71.64118 | *Pinus strobus* | MA |
| 008-01 | 7/20/01 | 42.18760 | -71.30645 | *Pinus strobus* | MA |
| 009-01 | 7/20/01 | 42.11954 | -71.32506 | *Pinus strobus* | MA |
| 011-01 | 7/22/01 | 42.51954 | -70.89672 | *Pinus strobus* | MA |
| 015-01 | 7/24/01 | 44.34897 | -68.34418 | *Pinus strobus* | ME |
| 108-04 | 7/18/04 | 35.8427167 | -86.42865 | *Pinus strobus* | TN |
| 110-04 | 7/18/04 | 36.1817667 | -86.301167 | *Pinus strobus* | TN |
| 112-04 | 7/19/04 | 35.9800333 | -85.015183 | *Pinus strobus* | TN |
| 121-04 | 7/19/04 | 35.7029 | -83.042067 | *Pinus strobus* | TN |
| 127-02 | 7/21/02 | 43.11453 | -71.09978 | *Pinus sylvestris* | NH |
| 129-04 | 7/22/04 | 37.1853833 | -81.128483 | *Pinus strobus* | VA |
| 130-02 | 7/21/02 | 43.11453 | -71.09978 | *Pinus sylvestris* | NH |
| 136-04 | 7/25/04 | 40.41585 | -74.444267 | *Pinus strobus* | NJ |
| 137-04 | 7/25/04 | 41.8812085 | -72.302025 | *Pinus strobus* | CT |
| 139-04 | 7/27/04 | 43.3236333 | -70.9888 | *Pinus strobus* | NH |
| 140-04 | 7/27/04 | 43.7135167 | -71.039383 | *Pinus resinosa* | NH |
| 145-02 | 8/8/02 | 45.92643 | -77.32558 | *Pinus banksiana* | ON |
| 146-04 | 7/27/04 | 43.7813333 | -71.170033 | *Pinus rigida* | NH |
| 150-02 | 8/8/02 | 45.92643 | -77.32558 | *Pinus banksiana* | ON |
| 150-04 | 7/27/04 | 43.8032333 | -70.748567 | *Pinus strobus* | ME |
| 152-02 | 8/8/02 | 45.93155 | -77.33315 | *Pinus strobus* | ON |
| 152-04 | 7/28/04 | 43.4008833 | -70.584683 | *Pinus strobus* | ME |
| 153-02 | 8/8/02 | 45.93155 | -77.33315 | *Pinus banksiana* | ON |
| 158-04 | 7/28/04 | 44.6220167 | -67.81875 | *Pinus strobus* | ME |
| 159-04A | 7/29/04 | 44.53105 | -68.3669 | *Pinus strobus* | ME |
| 166-04 | 7/30/04 | 44.01845 | -71.113533 | *Pinus strobus* | ME |
| 167-04 | 7/30/04 | 43.9599 | -71.191217 | *Pinus resinosa* | NH |
| 170-02 | 8/9/02 | 44.79382 | -78.07788 | *Pinus strobus* | ON |
| 175-04 | 8/14/04 | 43.939 | -90.623767 | *Pinus banksiana* | WI |
| 187-02 | 8/11/02 | 45.81847 | -80.54630 | *Pinus strobus* | ON |
| 187-04 | 8/14/04 | 44.4657333 | -91.636683 | *Pinus strobus* | WI |
| 194-04 | 8/15/04 | 43.91215 | -90.8659 | *Pinus banksiana* | MN |
| 200-04 | 8/15/04 | 44.0265667 | -89.809083 | *Pinus banksiana* | WI |
| 215-02 | 8/13/02 | 48.37632 | -89.58443 | *Pinus banksiana* | ON |
| 221-02 | 8/13/02 | 48.41557 | -89.63775 | *Pinus banksiana* | ON |
| 341-02 | 8/19/02 | 46.39492 | -79.24417 | *Pinus strobus* | ON |
| 346-02 | 8/19/02 | 46.39492 | -79.24417 | *Pinus strobus* | ON |
| AG034 | 6/24/19 | 39.3161 | -84.5723 | *Pinus strobus* | KY |
| AG036 | 6/25/19 | 39.6741 | -84.2271 | *Pinus strobus* | KY |

| | | | | | |
|---|---|---|---|---|---|
| AG042 | 6/28/19 | 40.3485 | -80.0187 | *Pinus strobus* | OH |
| AG043 | 6/28/19 | 39.872 | -82.9038 | *Pinus strobus* | OH |
| AG046 | 7/6/19 | 43.2105 | -87.8994 | *Pinus strobus* | WI |
| AG053 | 7/7/19 | 44.5066 | -87.9938 | *Pinus strobus* | WI |
| AG075 | 7/20/19 | 41.3525 | -72.1258 | *Pinus strobus* | CT |
| AG076 | 7/20/19 | 41.3452 | -72.1253 | *Pinus strobus* | CT |
| AG077 | 7/21/19 | 42.7553 | -71.2203 | *Pinus strobus* | CT |
| AG080 | 7/31/19 | 43.4255 | -89.4765 | *Pinus resinosa* | WI |
| AG081 | 7/31/19 | 43.751 | -89.9498 | *Pinus strobus* | WI |
| AG082 | 8/1/19 | 44.0628 | -92.4501 | *Pinus strobus* | MN |
| AG083 | 8/1/19 | 44.7572 | -93.1811 | *Pinus strobus* | MN |
| AG084 | 8/1/19 | 45.0653 | -93.1348 | *Pinus strobus* | MN |
| AG104 | 8/10/19 | 42.1659 | -85.5969 | *Pinus strobus* | MI |
| AG105 | 8/11/19 | 43.4595 | -85.4743 | *Pinus strobus* | MI |
| AG130 | 8/13/19 | 45.492 | -84.0294 | *Pinus strobus* | MI |
| AG138 | 8/12/19 | 45.5009 | -84.6146 | *Pinus strobus* | MI |
| AG140 | 8/21/19 | 38.4843 | -87.3034 | *Pinus strobus* | IL |
| CAN001 | 7/26/14 | 43.154604 | -80.675354 | *Pinus strobus* | MI |
| CAN061B | 8/1/14 | 43.789444 | -85.740278 | *Pinus banksiana* | MI |
| DS016 | 7/6/15 | 37.987968 | -84.532627 | *Pinus strobus* | KY |
| DS053 | 10/13/20 | 38.0409113 | -84.438831 | *Pinus strobus* | KY |
| DS055 | ?/?14 | 44.583074 | -90.38169 | *Pinus banksiana* | Wi |
| DS056 | 8/6/14 | 48.778368 | -88.55559 | *Pinus banksiana* | ON |
| DS057 | ?/?/14 | 45.124771 | -92.383617 | *Pinus banksiana* | WI |
| DS058 | 8/7/14 | 44.111957 | -84.120831 | *Pinus strobus* | MI |
| DS059 | 6/15/15 | 37.18483 | -81.130844 | *Pinus strobus* | VA |
| DS060 | 6/28/16 | 45.842776 | -91.887646 | *Pinus banksiana* | WI |
| DS061 | 6/28/16 | 45.1224 | -92.382533 | *Pinus strobus* | WI |
| DS062 | 6/27/15 | 46.311126 | -81.655729 | *Pinus banksiana* | ON |
| DS063 | 6/28/16 | 45.1224 | -92.382533 | *Pinus resinosa* | WI |
| DS064 | ?/?/14 | 44.858561 | -89.696097 | *Pinus banksiana* | WI |
| DS066 | 2014 | 45.124771 | -92.383612 | *Pinus resinosa* | WI |
| DS067 | 5/27/15 | 37.87584 | -79.308844 | *Pinus strobus* | VA |
| DS068 | 8/6/14 | 48.378056 | -89.580318 | *Pinus banksiana* | ON |
| DS069 | 6/13/15 | 35.442039 | -86.5277 | *Pinus taeda* | TN |
| DS071 | 8/17/15 | 46.508949 | -90.502659 | *Pinus banksiana* | WI |
| DS072 | 6/29/16 | 46.554426 | -91.322284 | *Pinus banksiana* | WI |
| DS073 | 6/28/15 | 45.966589 | -77.37125 | *Pinus banksiana* | ON |
| DS074 | 8/13/15 | 45.926308 | -77.325382 | *Pinus banksiana* | ON |
| DS075 | 6/27/15 | 46.311126 | -81.655729 | *Pinus sylvestris* | ON |
| LL016 | 7/16/13 | 37.0667 | -84.145833 | *Pinus strobus* | KY |
| RB109 | 8/3/12 | 41.05 | -77.150278 | *Pinus strobus* | PA |
| RB422 | 8/17/15 | 44.2260833 | -90.706756 | *Pinus banksiana* | WI |
| RB424 | 7/8/16 | 45.9178083 | -86.313008 | *Pinus resinosa* | MI |
| RB436 | 7/9/16 | 44.8438889 | -89.690556 | *Pinus resinosa* | WI |

**Table S2: Sample sequencing data, including total read count, how many total reads mapped to the reference genome, proportion of bases in the reference covered by at least one read, mean depth of reads at each base, and heterozygosity.**

| Sample name | Total read count | Reads mapped to reference | Reference coverage prop. | Mean base depth | Heterozygosity |
|---|---|---|---|---|---|
| 006-01 | 10007916 | 9502385 | 0.91875003 | 4.68230071 | 0.21698528 |
| 008-01 | 8838106 | 7204437 | 0.92106764 | 4.82829429 | 0.01757834 |
| 009-01 | 10502068 | 10108284 | 0.9323753 | 4.39755357 | 0.20423724 |
| 011-01 | 10957462 | 10535789 | 0.93578123 | 5.18390714 | 0.21720078 |
| 015-01 | 11478190 | 11045215 | 0.89915945 | 3.36356286 | 0.00599252 |
| 108-04 | 34642204 | 33097961 | 0.98062314 | 15.98965 | 0.18923904 |
| 110-04 | 15997902 | 15420899 | 0.93697118 | 5.40659214 | 0.20619308 |
| 112-04 | 11903290 | 11281022 | 0.86754524 | 2.85065071 | 0.0114684 |
| 121-04 | 12361598 | 11602827 | 0.89906231 | 3.17678786 | 0.08402471 |
| 127-02 | 53443148 | 50952916 | 0.88440838 | 3.30659286 | 0.00599929 |
| 129-04 | 13358294 | 12754052 | 0.89529238 | 3.17723714 | 0.01096205 |
| 130-02 | 30581038 | 29576899 | 0.97507161 | 11.0174957 | 0.10912428 |
| 136-04 | 13759830 | 13119342 | 0.93210586 | 5.329185 | 0.01464182 |
| 137-04 | 15545850 | 14945644 | 0.86710597 | 2.88306143 | 0.15287053 |
| 139-04 | 16182470 | 15336490 | 0.93898795 | 4.23258429 | 0.20736083 |
| 140-04 | 16457940 | 15814275 | 0.97273672 | 14.1046714 | 0.21991569 |
| 145-02 | 43725884 | 41761565 | 0.96366357 | 11.0682843 | 0.17101646 |
| 146-04 | 27635328 | 26624539 | 0.9771724 | 19.7330143 | 0.00823019 |
| 150-02 | 35111660 | 33440072 | 0.93511536 | 5.01412429 | 0.19799436 |
| 150-04 | 37962900 | 36577524 | 0.9418593 | 5.08463786 | 0.20257545 |
| 152-02 | 35319826 | 33977282 | 0.94015297 | 5.15714571 | 0.00839466 |
| 152-04 | 39146416 | 37713963 | 0.96837381 | 16.7965429 | 0.0105952 |
| 153-02 | 42342108 | 40780945 | 0.9693936 | 15.9156857 | 0.21096488 |
| 158-04 | 43122484 | 41516713 | 0.95901885 | 6.77148786 | 0.00783216 |
| 159-04A | 49851718 | 48058585 | 0.97305072 | 17.1136143 | 0.01052085 |
| 166-04 | 11402600 | 10937646 | 0.91614993 | 4.3047 | 0.18856213 |
| 167-04 | 13430446 | 12914934 | 0.93772837 | 5.74128786 | 0.00978136 |
| 170-02 | 44684400 | 42025004 | 0.96086921 | 15.3352143 | 0.17886923 |
| 175-04 | 14420674 | 13837196 | 0.95450626 | 8.21482071 | 0.14168665 |
| 187-02 | 47422588 | 45769103 | 0.95183405 | 11.1702721 | 0.22846603 |
| 187-04 | 22689294 | 21568762 | 0.97389086 | 17.0279643 | 0.00613661 |
| 194-04 | 25212340 | 24295471 | 0.97762904 | 17.0531357 | 0.21405004 |

| | | | | | |
|---|---|---|---|---|---|
| 200-04 | 27406966 | 26134794 | 0.98614733 | 19.1036429 | 0.20495448 |
| 215-02 | 27789236 | 26662534 | 0.94620254 | 9.08398071 | 0.19325798 |
| 221-02 | 35087254 | 33700968 | 0.93594033 | 4.46335857 | 0.00683165 |
| 341-02 | 38555430 | 36926287 | 0.93555024 | 5.36284286 | 0.20947449 |
| 346-02 | 53145694 | 51231298 | 0.97152457 | 21.7465429 | 0.17206266 |
| AG034 | 44747256 | 43085631 | 0.95889707 | 13.8094357 | 0.19171718 |
| AG036 | 9110468 | 7849446 | 0.97230332 | 10.2870964 | 0.19925861 |
| AG042 | 11597878 | 11129516 | 0.976943 | 12.1210686 | 0.18320788 |
| AG043 | 12639534 | 12097359 | 0.93537847 | 6.42074357 | 0.20042649 |
| AG046 | 41802698 | 40194756 | 0.96680549 | 8.41298786 | 0.18261309 |
| AG053 | 42441604 | 38601693 | 0.97638653 | 16.6979357 | 0.1333491 |
| AG075 | 8227942 | 7913697 | 0.9598512 | 6.13090071 | 0.21503242 |
| AG076 | 16846928 | 15393831 | 0.92658228 | 4.65414214 | 0.20110031 |
| AG077 | 17021266 | 15641980 | 0.90844044 | 5.000735 | 0.21205293 |
| AG080 | 17384360 | 16586279 | 0.94830642 | 6.379505 | 0.01357124 |
| AG081 | 20463888 | 19697604 | 0.95506324 | 14.0003929 | 0.1633804 |
| AG082 | 10734672 | 10196843 | 0.95738003 | 10.1483629 | 0.19548261 |
| AG083 | 8456624 | 7545709 | 0.93722052 | 4.65514357 | 0.17875723 |
| AG084 | 26366908 | 24705760 | 0.97101995 | 12.0576893 | 0.16531728 |
| AG104 | 31000686 | 29485226 | 0.97106591 | 18.1002929 | 0.00776339 |
| AG105 | 32353162 | 30527362 | 0.96722571 | 18.3517357 | 0.16913525 |
| AG130 | 39361990 | 33882327 | 0.95281815 | 5.13240357 | 0.02007387 |
| AG138 | 40828324 | 38999536 | 0.97808688 | 6.40014214 | 0.1413913 |
| AG140 | 9218944 | 8506872 | 0.93671137 | 6.19748 | 0.12443389 |
| CAN001 | 47269356 | 44179217 | 0.94562583 | 6.81838929 | 0.19937146 |
| CAN061 B | 13036510 | 12524195 | 0.97754881 | 15.00195 | 0.24681509 |
| DS016 | 8146346 | 7852994 | 0.9322728 | 3.94170357 | 0.12323896 |
| DS053 | 7448476 | 6889606 | 0.89497006 | 3.11278286 | 0.00674371 |
| DS055 | 9717060 | 8478165 | 0.92947223 | 4.50874929 | 0.16447891 |
| DS056 | 10463778 | 10061880 | 0.86446895 | 2.65925143 | 0.00789608 |
| DS057 | 11158182 | 10744708 | 0.98510816 | 17.0215714 | 0.22197784 |
| DS058 | 12784106 | 12205272 | 0.96076231 | 9.32053857 | 0.13759687 |
| DS059 | 13023406 | 12508338 | 0.96582545 | 8.87251071 | 0.13472453 |
| DS060 | 20860204 | 19756109 | 0.92421565 | 5.10441929 | 0.17707857 |
| DS061 | 21637656 | 20677174 | 0.91564626 | 4.20576 | 0.14353499 |
| DS062 | 22454138 | 21620372 | 0.95096558 | 5.19305786 | 0.18438743 |
| DS063 | 23233240 | 22248870 | 0.95555565 | 8.13582143 | 0.19520804 |
| DS064 | 26478626 | 25462707 | 0.97814342 | 10.532105 | 0.14069414 |

| | | | | | |
|---|---|---|---|---|---|
| DS066 | 32349806 | 31074679 | 0.98712891 | 17.2017071 | 0.00528129 |
| DS067 | 34800930 | 33576718 | 0.9772614 | 13.8663357 | 0.14314364 |
| DS068 | 42852140 | 41105430 | 0.97791447 | 12.4451071 | 0.20828735 |
| DS069 | 43130366 | 41605668 | 0.96379174 | 8.45708857 | 0.18368763 |
| DS071 | 10593870 | 10019672 | 0.9112939 | 4.130605 | 0.22990011 |
| DS072 | 12484296 | 12011465 | 0.93982998 | 4.91698071 | 0.16606996 |
| DS073 | 12554930 | 12053291 | 0.96207235 | 11.7341521 | 0.23282025 |
| DS074 | 12671686 | 12176228 | 0.96276063 | 13.8441571 | 0.19239273 |
| DS075 | 38492050 | 35689558 | 0.96745274 | 8.988985 | 0.18598433 |
| LL016 | 21876508 | 20860601 | 0.99222668 | 17.6271643 | 0.18259194 |
| RB109 | 12807712 | 12286305 | 0.96284811 | 16.1807929 | 0.23386914 |
| RB422 | 21898210 | 20589189 | 0.97205002 | 14.5961429 | 0.00639089 |
| RB424 | 13032942 | 12528702 | 0.96934578 | 8.81458357 | 0.20352978 |
| RB436 | 13591032 | 13075475 | 0.9465221 | 5.09167643 | 0.1889584 |

**Table S3: Number of scaffolds, contigs, and other basic metrics (L/N50, L/N90) representing the *D. similis* genome assembly.**

| | |
|---|---|
| Main genome scaffold total | 81 |
| Main genome contig total | 89 |
| Main genome scaffold sequence total | 270.226 MB |
| Main genome contig sequence total | 270.225 MB |
| Main genome scaffold L/N50 | 6/19.014 MB |
| Main genome contig L/N50 | 7/16.203 MB |
| Main genome scaffold L/N90 | 13/11.122 MB |
| Main genome contig L/N90 | 15/6.019 MB |
| Max scaffold length | 28.02 MB |
| Max contig length | 27.4 MB |
| Number of scaffolds > 50 KB | 28 |
| % main genome in scaffolds > 50 KB | 99.35% |

**Table S4: A summary of the output of the blobtools pipeline showing scaffold or contig length, coverage, coverage class, number of complete and duplicated BUSCOs, and taxanomic assignment based on local alignment (BLAST) to the NCBI nt and the Uniprot proteome databases.**

| # record | length | coverage | coverage class | complete BUSCOs | duplicated BUSCOs | species |
|---|---|---|---|---|---|---|
| NC_060105.1 | 18130230 | 90.0106 | medium | 344 | 1 | Neodiprion lecontei |
| NC_060106.1 | 22783663 | 96.0186 | medium | 527 | 2 | Neodiprion lecontei |
| NC_060107.1 | 19013855 | 92.1357 | medium | 495 | 12 | Neodiprion lecontei |
| NC_060108.1 | 27936105 | 84.3518 | medium | 445 | 1 | Neodiprion lecontei |
| NC_060109.1 | 10882974 | 69.7112 | medium | 50 | 0 | Neodiprion lecontei |
| NC_060110.1 | 25438219 | 96.9604 | medium | 625 | 1 | Neodiprion lecontei |
| NC_060111.1 | 11122181 | 81.4412 | medium | 210 | 3 | Neodiprion lecontei |
| NC_060112.1 | 28020057 | 94.9402 | medium | 791 | 7 | Neodiprion lecontei |
| NC_060113.1 | 14403014 | 76.3266 | medium | 216 | 4 | Neodiprion lecontei |
| NC_060114.1 | 22560868 | 90.2777 | medium | 562 | 8 | Neodiprion lecontei |
| NC_060115.1 | 16102462 | 81.013 | medium | 315 | 3 | Neodiprion lecontei |
| NC_060116.1 | 17816656 | 88.9547 | medium | 386 | 4 | Neodiprion lecontei |
| NC_060117.1 | 16048524 | 71.9859 | medium | 224 | 3 | Neodiprion lecontei |
| NC_060118.1 | 14306973 | 80.1617 | medium | 267 | 3 | Neodiprion lecontei |
| NW_025724968.1 | 125000 | 9.491 | low | 0 | 0 | Diprion pini |
| NW_025724969.1 | 250000 | 57.8107 | medium | 0 | 0 | Diprion pini |
| NW_025724970.1 | 125000 | 8.2963 | low | 0 | 0 | Diprion pini |
| NW_025724971.1 | 38377 | 268.7409 | high | 0 | 0 | Monoctenus juniperi |
| NW_025724972.1 | 2657045 | 91.2721 | medium | 0 | 0 | Diprion pini |
| NW_025724973.1 | 63067 | 53.6455 | medium | 0 | 0 | Monoctenus juniperi |
| NW_025724974.1 | 67718 | 135.5745 | medium | 0 | 0 | Monoctenus juniperi |
| NW_025724975.1 | 95342 | 98.4956 | medium | 0 | 0 | Monoctenus juniperi |
| NW_025724976.1 | 34974 | 5.689 | low | 0 | 0 | Monoctenus juniperi |
| NW_025724977.1 | 49256 | 24.7718 | low | 0 | 0 | Monoctenus juniperi |
| NW_025724978.1 | 51213 | 51.3241 | medium | 0 | 0 | Monoctenus juniperi |
| NW_025724979.1 | 42338 | 475.1521 | high | 0 | 0 | Monoctenus juniperi |
| NW_025724980.1 | 26228 | 65.2935 | medium | 0 | 0 | Monoctenus juniperi |
| NW_025724981.1 | 40330 | 40.5411 | medium | 0 | 0 | Monoctenus juniperi |
| NW_025724982.1 | 25685 | 49.8144 | medium | 0 | 0 | Monoctenus juniperi |
| NW_025724983.1 | 51375 | 245.2921 | high | 0 | 0 | Monoctenus juniperi |
| NW_025724984.1 | 39764 | 389.5047 | high | 0 | 0 | Monoctenus juniperi |
| NW_025724985.1 | 28663 | 224.0174 | medium | 0 | 0 | Monoctenus juniperi |
| NW_025724986.1 | 158925 | 49.4756 | medium | 0 | 0 | Diprion pini |
| NW_025724987.1 | 33276 | 51.4669 | medium | 0 | 0 | Monoctenus juniperi |

| | | | | | | |
|---|---|---|---|---|---|---|
| NW_025724988.1 | 32071 | 40.4626 | medium | 0 | 0 | Monoctenus juniperi |
| NW_025724989.1 | 11842 | 6.192 | low | 0 | 0 | Asiemphytus rufocephalus |
| NW_025724990.1 | 85647 | 93.2507 | medium | 0 | 0 | Monoctenus juniperi |
| NW_025724991.1 | 33819 | 102.0365 | medium | 0 | 0 | Monoctenus juniperi |
| NW_025724992.1 | 35164 | 77.4848 | medium | 0 | 0 | Monoctenus juniperi |
| NW_025724993.1 | 67013 | 353.7247 | high | 0 | 0 | Monoctenus juniperi |
| NW_025724994.1 | 21015 | 19.5222 | low | 0 | 0 | Monoctenus juniperi |
| NW_025724995.1 | 25556 | 44.3266 | medium | 0 | 0 | Hoplocampa fulvicornis |
| NW_025724996.1 | 31078 | 22.0725 | low | 0 | 0 | Monoctenus juniperi |
| NW_025724997.1 | 27476 | 96.613 | medium | 0 | 0 | Monoctenus juniperi |
| NW_025724998.1 | 29750 | 8.3657 | low | 0 | 0 | Monoctenus juniperi |
| NW_025724999.1 | 30803 | 15.0811 | low | 0 | 0 | Monoctenus juniperi |
| NW_025725000.1 | 40699 | 17.0122 | low | 0 | 0 | Monoctenus juniperi |
| NW_025725001.1 | 25590 | 24.0322 | low | 0 | 0 | Monoctenus juniperi |
| NW_025725002.1 | 17887 | 105.883 | medium | 0 | 0 | Monoctenus juniperi |
| NW_025725003.1 | 23739 | 24.9112 | low | 0 | 0 | Neodiprion sp. BYU_ACHY028 |
| NW_025725004.1 | 32873 | 138.0812 | medium | 0 | 0 | Monoctenus juniperi |
| NW_025725005.1 | 24265 | 55.7459 | medium | 0 | 0 | Monoctenus juniperi |
| NW_025725006.1 | 34332 | 40.2138 | medium | 0 | 0 | Monoctenus juniperi |
| NW_025725007.1 | 42121 | 187.9828 | medium | 0 | 0 | Monoctenus juniperi |
| NW_025725008.1 | 25279 | 755.0142 | high | 0 | 0 | Liriomyza huidobrensis |
| NW_025725009.1 | 37521 | 45.6505 | medium | 0 | 0 | Monoctenus juniperi |
| NW_025725010.1 | 46645 | 12.7399 | low | 0 | 0 | Monoctenus juniperi |
| NW_025725011.1 | 44291 | 12.1009 | low | 0 | 0 | Monoctenus juniperi |
| NW_025725012.1 | 49816 | 34.3112 | medium | 0 | 0 | Monoctenus juniperi |
| NW_025725013.1 | 29950 | 75.0183 | medium | 0 | 0 | Monoctenus juniperi |
| NW_025725014.1 | 27887 | 10.6356 | low | 0 | 0 | Monoctenus juniperi |
| NW_025725015.1 | 38840 | 66.2661 | medium | 0 | 0 | Monoctenus juniperi |
| NW_025725016.1 | 42685 | 5.3754 | low | 0 | 0 | Monoctenus juniperi |
| NW_025725017.1 | 32062 | 46.7711 | medium | 0 | 0 | Monoctenus juniperi |
| NW_025725018.1 | 31230 | 24.7925 | low | 0 | 0 | Monoctenus juniperi |
| NW_025725019.1 | 33651 | 33.8218 | medium | 0 | 0 | Monoctenus juniperi |
| NW_025725020.1 | 35221 | 59.5464 | medium | 0 | 0 | Monoctenus juniperi |
| NW_025725021.1 | 27305 | 110.7911 | medium | 0 | 0 | Monoctenus juniperi |
| NW_025725022.1 | 50777 | 75.1312 | medium | 0 | 0 | Monoctenus juniperi |
| NW_025725023.1 | 68117 | 161.7321 | medium | 0 | 0 | Monoctenus juniperi |

| | | | | | | |
|---|---|---|---|---|---|---|
| NW_025725024.1 | 27312 | 111.4197 | medium | 0 | 0 | Monoctenus juniperi |
| NW_025725025.1 | 30247 | 54.952 | medium | 0 | 0 | Monoctenus juniperi |
| NW_025725026.1 | 28995 | 52.4401 | medium | 0 | 0 | Monoctenus juniperi |
| NW_025725027.1 | 41302 | 25.7811 | low | 0 | 0 | Monoctenus juniperi |
| NW_025725028.1 | 30455 | 43.9349 | medium | 0 | 0 | Monoctenus juniperi |
| NW_025725029.1 | 42670 | 43.5863 | medium | 0 | 0 | Monoctenus juniperi |
| NW_025725030.1 | 40155 | 72.0923 | medium | 0 | 0 | Monoctenus juniperi |
| NW_025725031.1 | 24401 | 38.4178 | medium | 0 | 0 | Monoctenus juniperi |
| NW_025725032.1 | 34746 | 83.778 | medium | 0 | 0 | Monoctenus juniperi |
| NW_025725033.1 | 33915 | 74.5624 | medium | 0 | 0 | Monoctenus juniperi |
| NW_025725034.1 | 28164 | 21.8174 | low | 0 | 0 | Monoctenus juniperi |

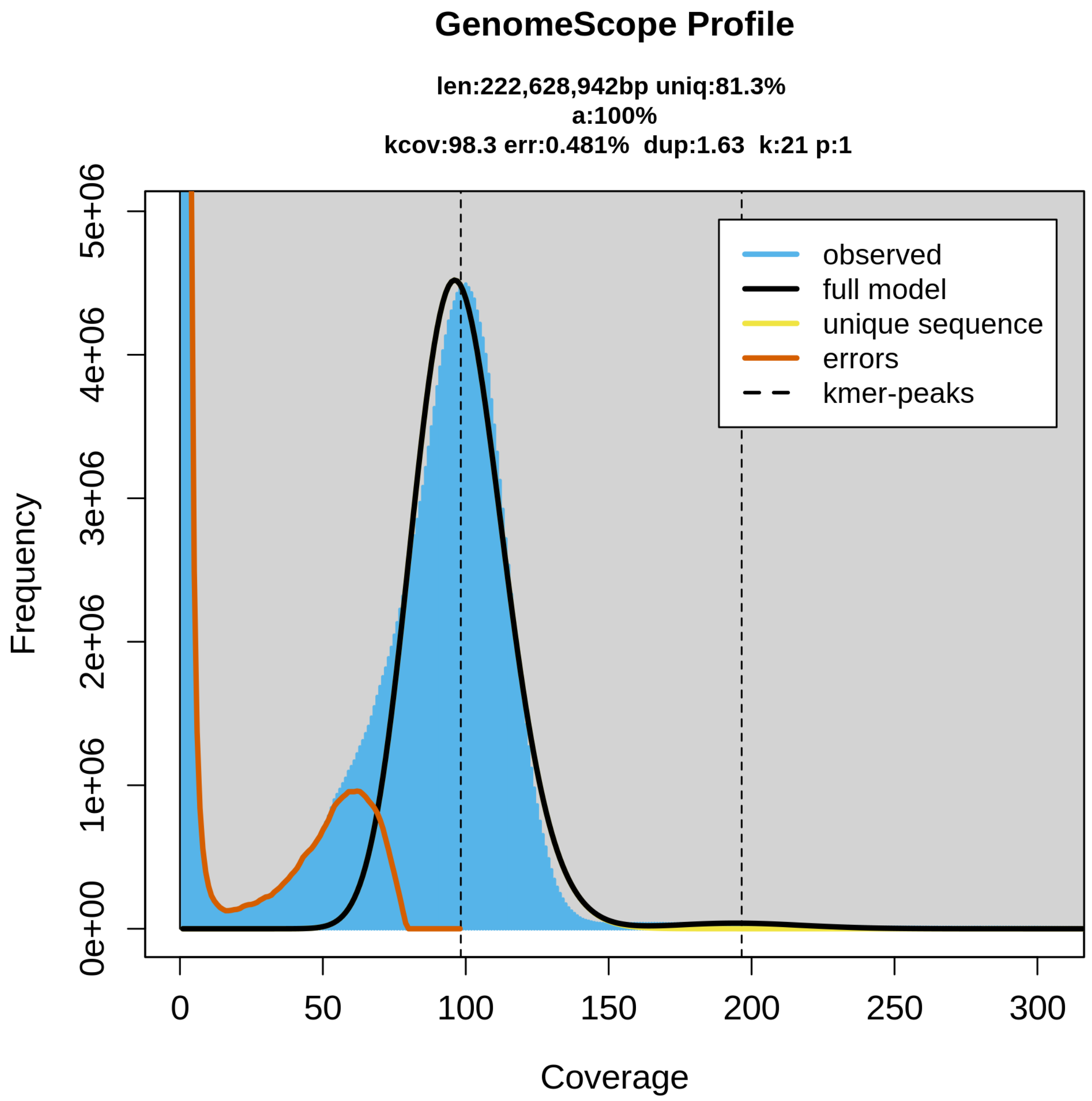


**Figure S1: K-mer profile output by GenomeScope2. Using a k-mer profile determine by KMC from the raw HiFi reads, GenomeScope2 applies a mathematical model to the k-mer frequencies to estimate heterozygosity, repeat content, and genome size depending on the known ploidy of the individual from which the reads were derived.**

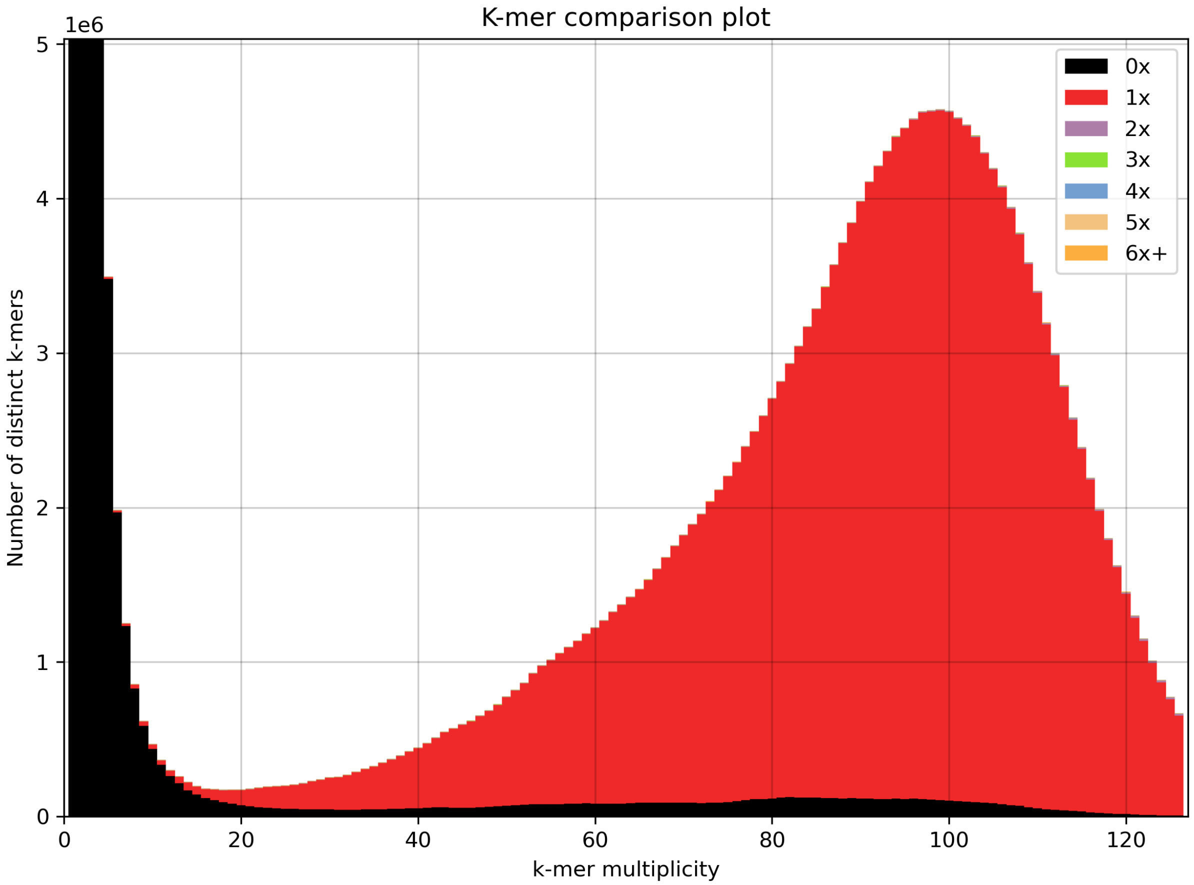


**Figure S2: Analysis of the genome assembly using k-mer spectra. K-mer spectra analysis of the raw reads used to generate the assembly and the assembly itself reveals the representation and abudance (multiplicity) of all kmers in the assembly in the x-axis. The k-mers represented in the black bars are k-mers in the raw read dataset that are not included in the assembly, and the k-mers represented in the red bars are in the assembly once which is expected in a haploid assembly. The dominant peak represents the average coverage of the genome as the largest number of k-mers are at that multiplicity. The absence of a second peak at half the multiplicity of the dominant peak indicates that there is no heterozygosity in this sample, which is expected of a haploid male.**

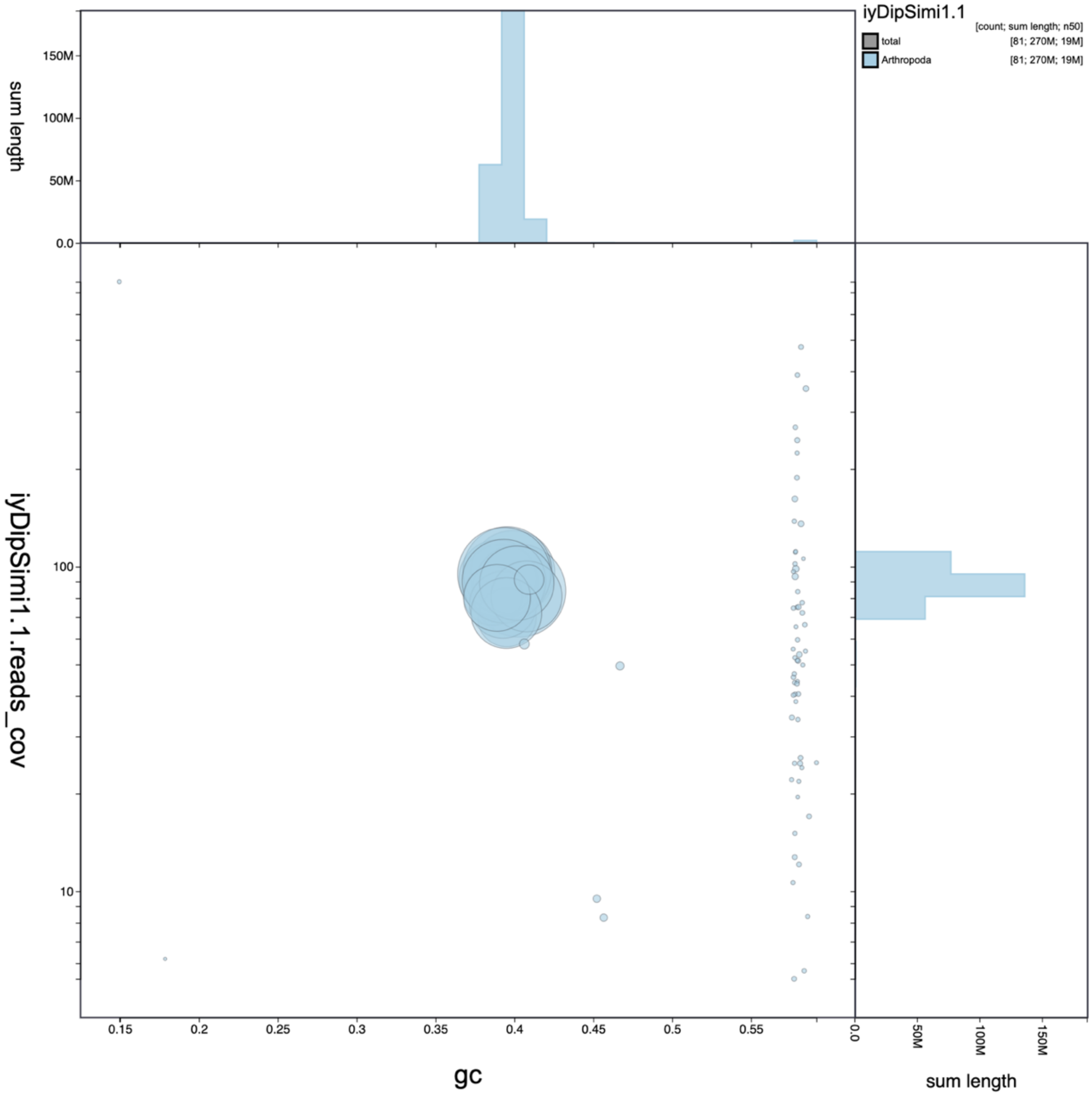


**Figure S3: This is an output of blobtookit which shows GC content (x-axis) read coverage (y-axis) and taxonomic identity of the fragments in the genome. The clustering of fragments assigned to certain taxa combined with read coverage, fragment length, and GC content were used to assess for the presence of contaminant contigs in the assembly.**

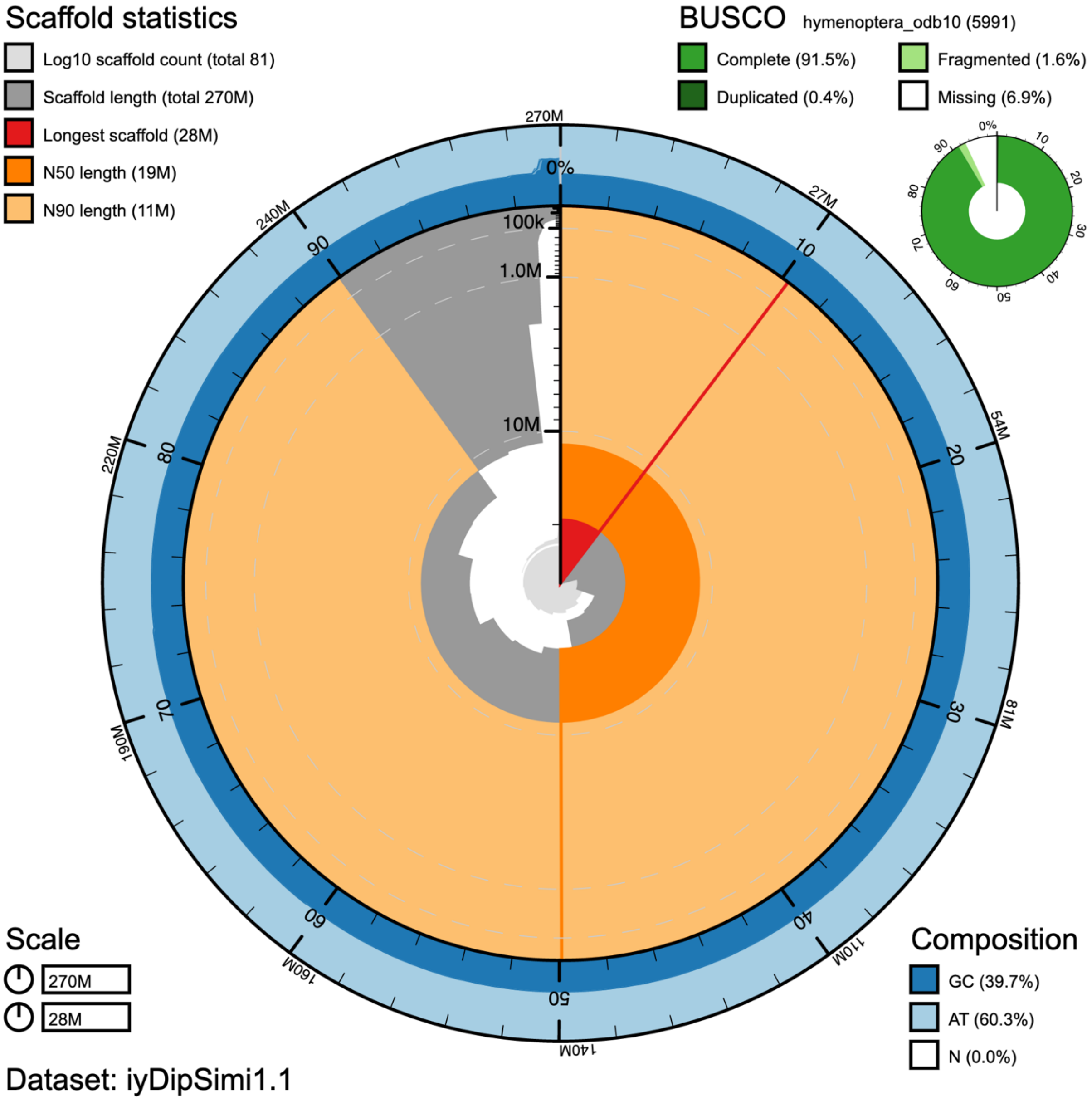


**Figure S4: Snail plot of the final scaffolded assembly. A product of the blobtoolkit, this represents the genome as a circle. The circumference represents the size of the genome (270 MB), and the radius represents the length of the longest fragment (28 Mb). The scaffolds are ordered from largest to smallest starting at the top of the circle going clockwise. The dark orange represents the scaffold N50 (19 Mb) and the light orange represents the scaffold N90 (11 Mb). The results of the BUSCO analysis using the Hymenoptera ortholog dataset v.10 is depicted in the top right, and GC content for every fragment is represented by the dark blue bar around the perimeter of the circle with an average of 39.7%.**

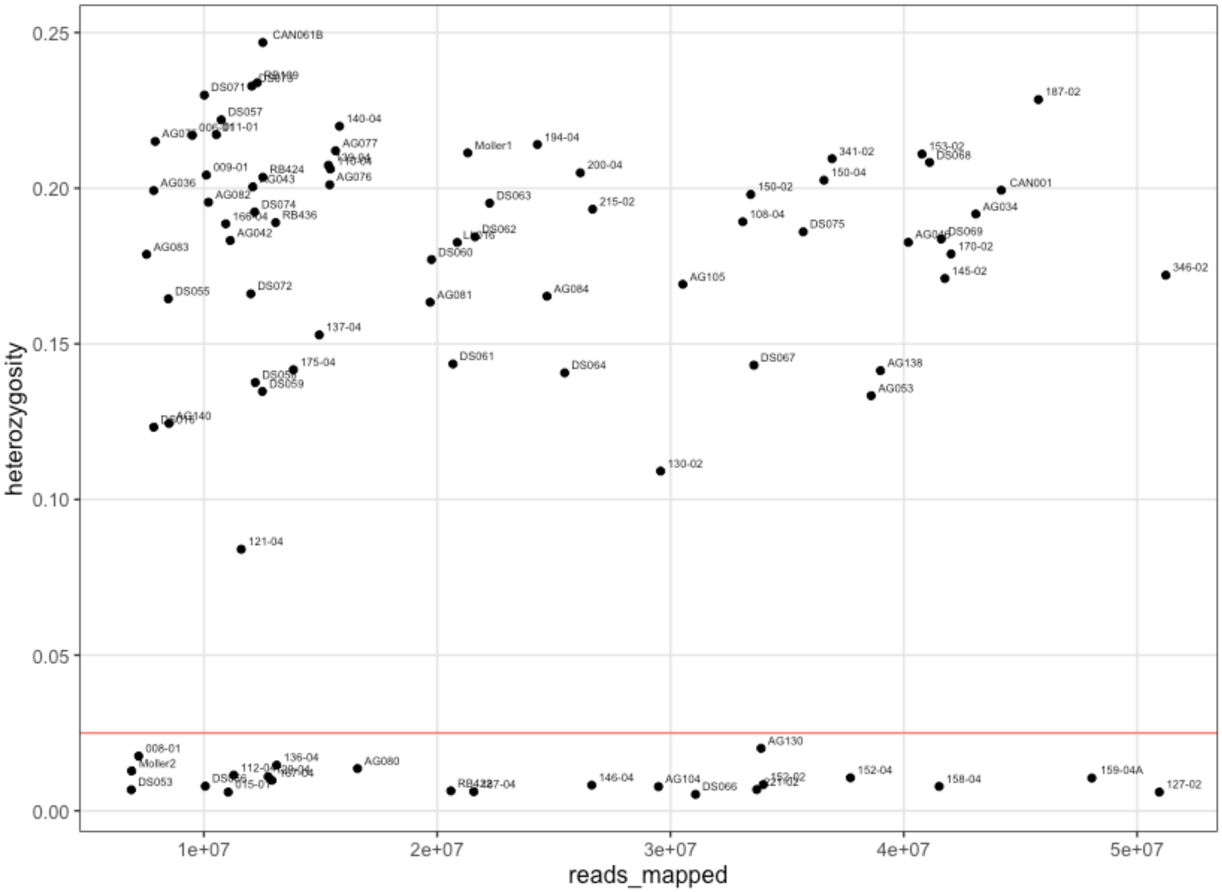


**Figure S5: Total number of reads mapped to genome and heterozygosity for all 84 sequenced samples. Red horizontal line at 0.025 heterozygosity was the threshold used to determine sex and ploidy of samples, with samples below this threshold deemed haploid males and excluded from further population analyses. Note the stark contrast between groups.**

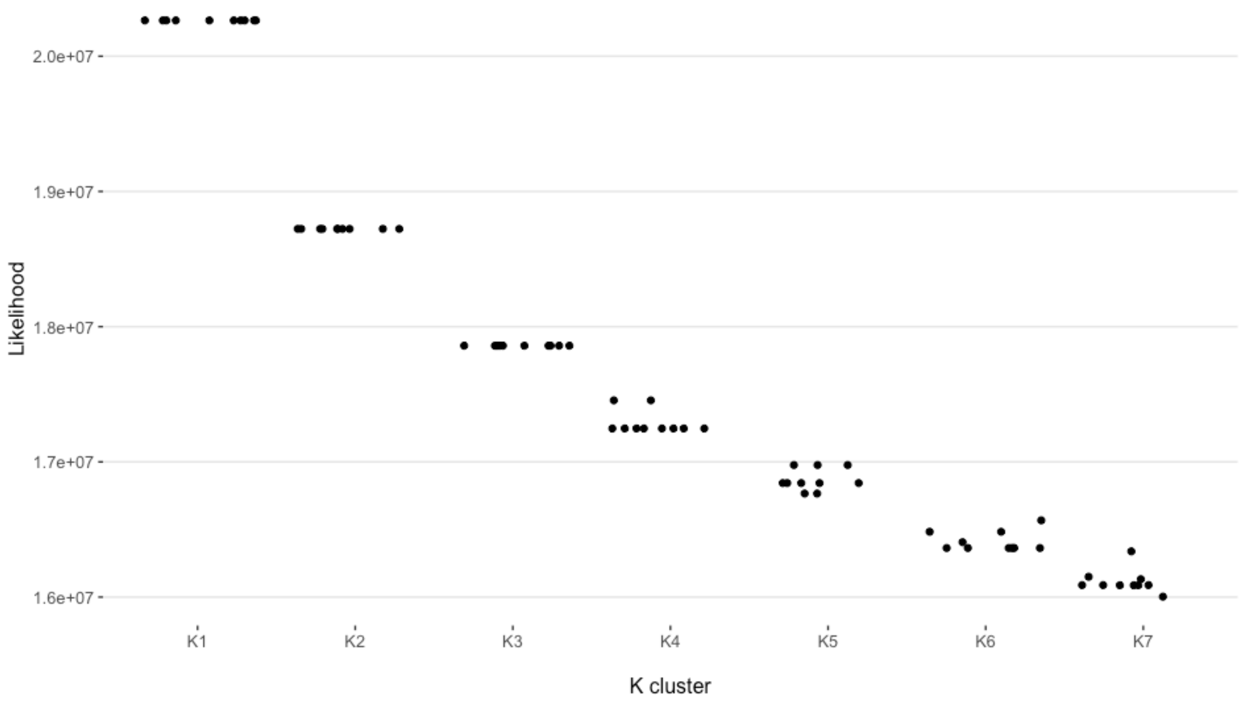


**Figure S6: Likelihood results from 10 replicates of NgsAdmix run at each K cluster from 1-7. Resulting likelihoods for K=1,2,3 were identical for each run. These results show a clear cascading likelihood of a single genetic cluster as the most likely genetic grouping.**

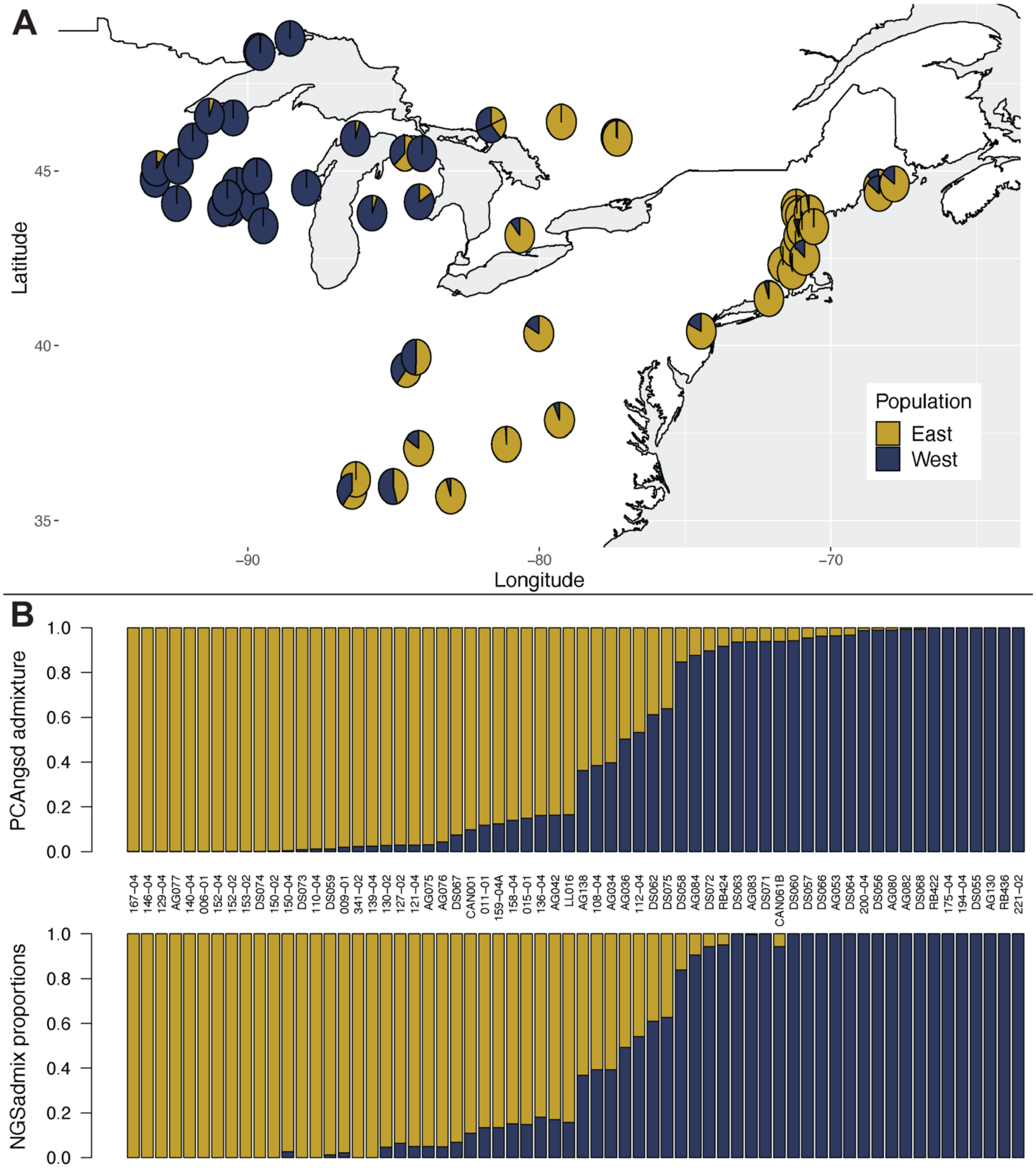


**Figure S7: Admixture proportions and collection locations for each of the 64 female *D. similis* samples used in this study, showing East and West subpopulation ancestry in gold and indigo respectively. Panel A shows collection sites and admixture as determined by PCAngsd analysis using *K* = 2 (see methods). Panel B shows admixture proportions output for *K* = 2 from two programs: PCAngsd on top (the same as displayed on the map), and NgsAdmix on the bottom**
